# The tumor necrosis superfamily member 4-1BBL expressed by tumor cells prevents exhaustion of CD8 T cells in a humanized mouse model of papillary renal cell carcinoma

**DOI:** 10.1101/2025.07.25.666825

**Authors:** Marie Fornier, Julien Novarino, Marie Naturel, Marylou Panouillot, Marie-Caroline Dieu-Nosjean, Gilles Marodon

## Abstract

**Background:** 4-1BB (CD137), a member of the TNF receptor superfamily, is a critical co-stimulatory receptor for CD8⁺ T cell activation and regulatory T cell (Treg) expansion. While its ligand 4-1BBL is typically expressed by professional antigen-presenting cells, several carcinomas also express 4-1BBL, though its function in the tumor microenvironment remains poorly defined.

**Methods:** We analyzed 4-1BBL expression across human tumors and found papillary renal cell carcinoma (pRCC) to exhibit the highest levels. Using The Cancer Genome Atlas, we found high 4-1BBL expression correlated with poor overall survival in pRCC. To study its role in vivo, we established an orthotopic humanized mouse model of pRCC by grafting ACHN cells into the renal capsule of mice reconstituted with human CD34⁺ hematopoietic stem cells. We then performed CRISPR-mediated deletion of 4-1BBL in tumor cells, followed by flow cytometry and single-cell RNA sequencing of tumor-infiltrating immune cells.

**Results:** Loss of tumor-derived 4-1BBL resulted in accelerated tumor growth and decreased immune cell clustering. In the absence of 4-1BBL, CD8⁺ T cells displayed elevated expression of PD-1, TIM-3, LAG-3, granzyme B, perforin, and NKG7, indicating a cytotoxic yet exhausted phenotype. Treg were only modestly impacted. Tumor-infiltrating CD8⁺ T cells expressed high levels of 4-1BBL and showed transcriptional signatures of altered AP-1 factors and enhanced PI3K pathway signaling.

**Conclusions:** Our findings uncover a previously unrecognized role for tumor- and T cell–derived 4-1BBL in sustaining cytotoxic CD8⁺ T cell functionality and limiting their exhaustion. This reveals a potential immune-regulatory axis that could be exploited for therapeutic modulation in renal cell carcinoma.

## Introduction

The immune response to tumors is orchestrated by a complex network of cells and molecules, resulting in either tolerance or rejection. Although limited to a small number of patients, the efficacy of immune checkpoint inhibitors (ICI) proves that the immune system is capable of rejecting tumors and curing cancer. To transpose this principle into therapy, more basic research on co-stimulatory molecules and tolerance mechanisms to tumors is needed. Among those mechanisms, inhibitory signals directly delivered to T cells, such as PD-1, appear crucial. Other immunosuppressive mechanisms are more indirect, such as the action of regulatory T cells (Treg) on the immune response to tumors. Indeed, Tregs are often correlated to bad prognosis in various cancers (1) and countless studies in mice have proven the efficacy at depleting Treg (2). It should be noted however that the efficacy of this strategy in cancer patients remains to be proven (3).

As stated above, intratumoral T cells interact with antigen presenting cells (APC), other immune cells and the tumor itself (notwithstanding the nerves), the result of this temporal dialog being either rejection (tumor elimination) or immune repression (tumor growth). The nature of this dialog is governed by chemokines and cytokines but also by direct cell contact through the T cell (antigen) receptor and various co stimulatory molecules with activating (such as CD28, ICOS, or 4-1BB) or inhibiting (such as PD-1,TIGIT, TIM3, LAG3, or BTLA) properties. It is important to stress that Treg often express the highest levels of those co-stimulatory and co-inhibitory receptors in the tumor micro-environment (TME), raising concerns about the use of co-stimulatory agonists that may also target Treg and favor immunosuppression (4). Those co-stimulatory molecules possess a set of ligands that are normally expressed by immune cells, especially APCs, but that can also be aberrantly expressed by cancer cells themselves. Moreover, cancer cells often express MHC class I and/or class II molecules which likely impacts the CD8 and CD4 T cell response, including Treg (5). Although the interactions between T cells and APCs are primordial, the dialog between T cells and cancer cells undoubtedly participates in the outcome of the immune response to tumors.

Among the many co-stimulatory pairs expressed by tumors, 4-1BB (CD137, encoded by the *TNFRSF9* gene) has attracted much attention due to the high efficacy of agonist antibodies in syngeneic models (6). Unfortunately, these positive results with 4-1BB agonists did not translate into humans for various reasons (7). Nevertheless, the high potential of 4-1BB for T cell activation has led to the inclusion of the 4-1BB signaling domain into CAR-T cells to favor their persistence and function (8). In addition, this has been shown to prevent exhaustion of CAR-T cells due to tonic signaling (9). Signaling through 4-1BB is mostly TRAF-dependent with subsequent NF-kB and ERK activation, which protects cells from apoptosis and DNA damage (10).

Numerous publications have shown that 4-1BB is indeed a major co-stimulatory molecule for full blown effector CD8 T cell-mediated immune response (11). In addition, 4-1BB was recently shown to be a major inducer of proliferation of exhausted CD8 T cells and terminal differentiation with positive outcome on tumor growth in murine syngeneic models (12). Aside from this proven role on CD8 T cells, there is also a consensus that 4-1BB is a conserved intratumoral Treg marker (13,14). This has led to the demonstration that depleting Treg through 4-1BB Fc-mediated depletion (ADCC) leads to better tumor control (15). Our own results have shown that targeting 4-1BB on murine Treg in vitro deeply alters their transcriptome profile and enhances their suppressive function in vivo (16). Others have shown that 4-1BB on Treg is used to strip 4-1BBL from the APC cell surface (trogocytosis), preventing the latest to fully activate neighboring T cells (17). The unique ligand of 4-1BB, 4-1BBL (CD137L, encoded by the *TNFSF9* gene), is essential for 4-1BB signaling in T cells. This is most often performed by professional antigen presenting cells, such as dendritic cells or B cells but expression of 4-1BBL has also been reported in various carcinomas (18). The functional consequence of that expression on CD8 T cells or on Treg in a cancer context is unclear.

A recent publication reports that expression of 4-1BBL on extracellular vesicles is able to generate Treg with functional consequences on the growth of the tumor in a syngeneic model (19). To add to the complexity, 4-1BBL is able to bind to 4-1BB on the same cell to induce *cis*-activation (20,21). Moreover, 4-1BBL is also able to transmit reverse signaling upon binding with 4-1BB with various consequences depending on the affected cell (11). Thus, in a cancer immunotherapy context, 4-1BB agonists might activate CD8 effector and regulatory T cells alike, and given the numerous consequences that the agonists might have on CD8 or on Treg, functional outcomes of the treatment would be difficult to predict. Thus, we believe that the natural biological function of 4-1BB co-stimulation on T cells in situ would be best appreciated in conditions where the T cell immune response is developing in the absence of the ligand, rather than with 4-1BB agonists that may induce unforeseen phenotypes and cellular functions. In that respect, it has been clearly established that the two 4-1BB agonists available in the clinic have different abilities to block the binding of 4-1BBL to 4-1BB or to prevent trimerization of the ligand (22,23).

It is also important to realize that the murine 4-1BB is only 60% homologous to its human counterpart. Moreover, human 4-1BBL can bind to mouse 4-1BB, but mouse 4-1BBL cannot bind to human 4-1BB, and the combining affinity of human 4-1BBL to mouse 4-1BB is about 30% of that of human 4-1BB (24). Thus, syngeneic models might poorly reflect the 4-1BB/4-1BBL biology in humans. To circumvent these limitations, we leveraged mice reconstituted with a human immune system to decipher the dialog between a renal cancer cell line and T cells in vivo. We chose kidney cancer as a model since analysis of the TCGA dataset shows that 4-1BBL expression negatively correlates with overall survival in kidney cancer patients. Thus, using a 4-1BBL-deficient kidney cancer cell line, multi parameter flow cytometry and single cell RNA sequencing, we demonstrate that 4-1BBL expressed by tumor cells exerts limited influence on Treg biology, but is a major regulator of T cell exhaustion.

## Results

### 4-1BBL is a biomarker for poor prognosis in papillary kidney cancer patients

To investigate the role of 4-1BBL expression in the immune response to cancer, we first identified which cancer types exhibit the highest levels of 4-1BBL expression. Analysis of the Human Protein Atlas reveals that the expression of 4-1BBL is the highest in kidney cancer cell lines (supplemental figure 1A). In agreement with this observation, the expression of 4-1BBL in clear cell (ccRCC) and papillary (pRCC) renal cell carcinomas, the two main subtypes of kidney cancer, is the highest of all solid tumors of the TCGA dataset (supplemental figure 1B). Moreover, 4-1BBL expression is much higher in the tumor than in the normal tissue for pRCC and ccRCC (supplemental figure 1C).

Analysis of the TCGA dataset shows that overall survival of patients is negatively associated with the expression level of 4-1BBL for pRCC but not ccRCC (supplemental figure 2A). In the same line, 4-1BBL expression correlates with advancement of the disease (clinical stages) for pRCC and ccRCC (supplementary figure 2B). The same trend is also observed for Pancreatic adenocarcinoma (PAAD), Bladder urothelial carcinoma (BLCA), Mesothelioma (MESO) and Uveal melanoma (UVM) (supplementary figure 2C), showing that 4-1BBL is a bad prognosis factor in several cancers in addition to pRCC.

### An orthotopic model of papillary RCC in immunodeficient mice

To investigate further the biological function of 4-1BBL in kidney cancer, we aim to develop a humanized mice model of pRCC, possibly more relevant to human physiology than syngeneic mouse models. As proof-of-principle, we chose the ACHN cell line since it is genetically close to tumors of pRCC patients (25). We first show that the ACHN cell line expresses 4-1BBL at both the mRNA and protein levels (Figure 1A). Thus, we grafted 2.10^6^ cells under the renal capsule of immunodeficient NXG mice and observed the anatomical structures of the tumor at the end of the experiment (Figure 1B). Histological examination of the kidney (zone A) revealed distinct morphological features across three representative regions. In the peripheral area (zone B), the tumor displayed a sharp interface with adjacent normal kidney parenchyma and was composed of epithelial cells arranged in tubular and short papillary structures embedded in a delicate fibrovascular stroma, consistent with pRCC morphology. In a more central area (zone C), extensive coagulative necrosis was present, bordered by sheets of viable tumor cells, indicative of aggressive tumor behavior and poor vascular supply. In contrast, high-magnification analysis of a third region (zone D) revealed a fascicular architecture composed of spindle-shaped cells with elongated nuclei, absent papillary or tubular structures, consistent with a sarcomatoid component. These features collectively support the classification of the ACHN-derived tumor as a high-grade pRCC with sarcomatoid differentiation.

**Figure 1.**
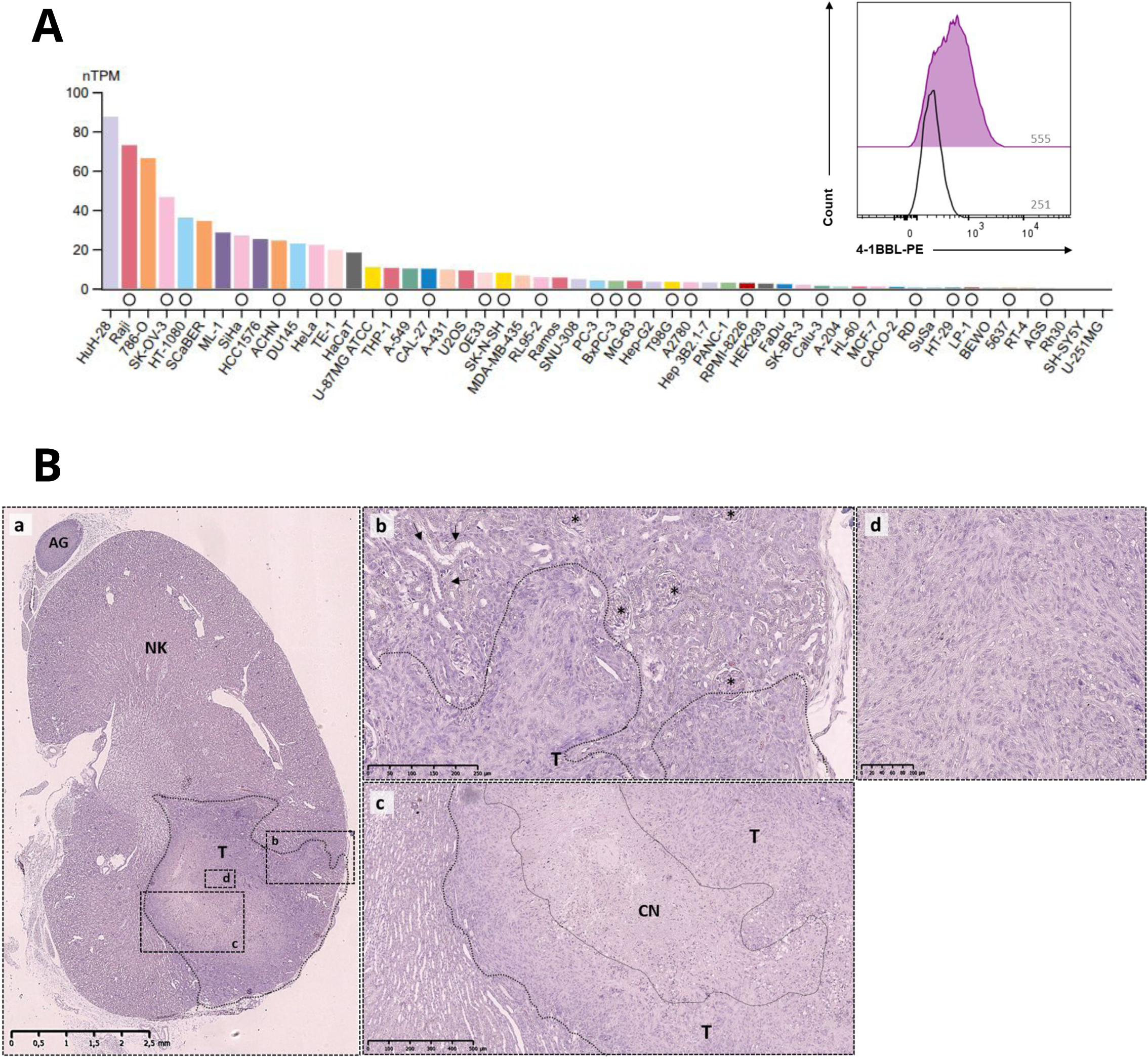
An orthotopic model of papillary RCC in immunodeficient mice. **(A)** Transcriptomic analysis of *TNFSF9* (4-1BBL) expression across a panel of human cancer cell lines (The Protein Altas). Expression is shown in normalized transcripts per million (nTPM). Inset: flow cytometry histogram showing surface 4-1BBL protein levels in ACHN cell line (purple) compared to isotype control (black). Numbers indicate mean fluorescence intensity (MFI). **(B)** Histopathological features of a renal tumor grafted in NXG mice **(a)** Low-magnification overview of the kidney showing a well-demarcated tumor (T) compressing the normal parenchyma (NK) and sparing the adrenal gland (AG). The dotted line delineates the tumor boundary. (scale bar unit: 0.5 mm) **(b)** Higher magnification of the tumor periphery (boxed region b in (a)) reveals epithelial cells arranged in tubular and short papillary structures (asterisks), consistent with papillary carcinoma morphology. Arrowheads indicate the boundary with adjacent normal tubules. (scale bar unit: 50 μm) **(c)** Intermediate magnification of a central tumor region (boxed region c in (A)), showing extensive central necrosis (CN) surrounded by viable tumor tissue (T). (scale bar unit: 100 μm) **(d)** High-magnification image of a sarcomatoid area (boxed region d in (A)), (scale bar unit: 20 μm).

### 4-1BBL-deficiency does not impact growth of a pRCC cancer cell line in immunodeficient hosts

Then, we wanted to investigate the role of tumoral 4-1BBL expression on the immune response to pRCC *in vivo*. Thus, we generated an ACHN kidney cancer cell line in which the TNFSF9 gene was “crispered” out to permanently abolish expression of 4-1BBL. The subcloning step necessary to isolate a single mutant introduces a bias due to selection of fastest growing clones. To be sure that we compare cell lines differing only by 4-1BBL expression, we restored expression of 4-1BBL in the deficient clone with a lentivirus encoding the TNFSF9 gene and the GFP reporter gene (Lv-pCMV-TNFSF9-GFP). As a control, the same deficient clone was transduced with a lentivirus expressing the GFP reporter gene alone. As expected, all GFP+ cells express 4-1BBL with the Lv-pCMV-TNFSF9-GFP vector whereas none express 4-1BBL with the control Lv–pCMV-GFP vector (Figure 2A). To monitor tumor growth *in vivo*, a second transduction with a mCherry/luciferase-encoding lentiviral vector was performed in the bulk populations of the two cell lines (Figure 2B). The resulting cell lines were then implanted in the renal capsule of immunodeficient NXG mice, and tumor growth was monitored by bioluminescence quantification (Figure 2C). We observe that tumor growth is similar in the two conditions (Figure 2D), indicating that the 4-1BBL status of the tumor does not impact viability and growth in immunodeficient hosts.

**Figure 2.**
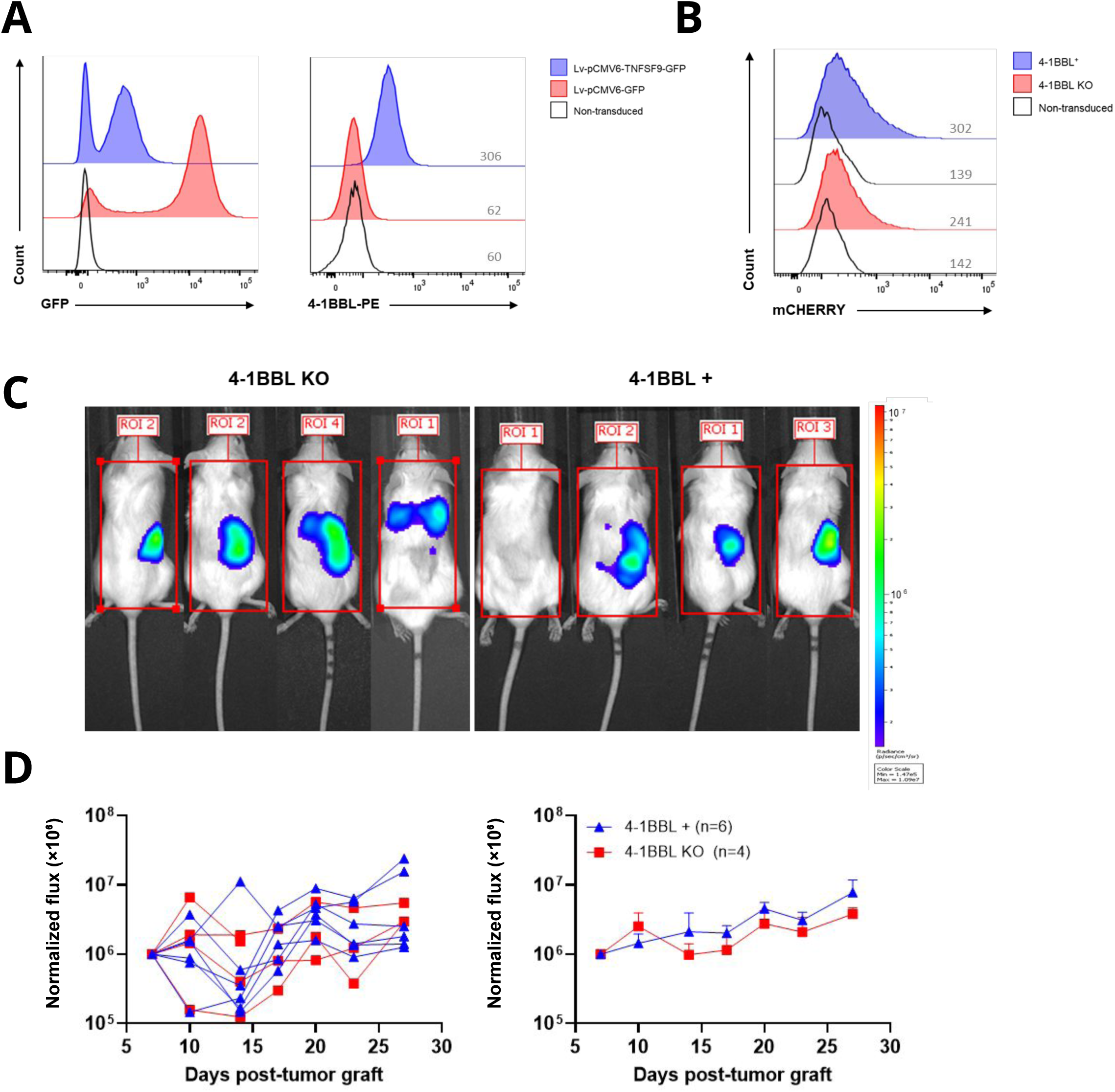
4-1BBL-deficiency does not impact growth of a pRCC cancer cell line in immunodeficient hosts. **(A)** Flow cytometry validation of ACHN tumor cells transduced with lentiviral constructs encoding either *TNFSF9*-GFP (blue), GFP alone (red) or non-transduced cells (black). Left panel: GFP expression 14 days after transduction; right panel: surface expression of 4-1BBL (PE) in GFP+ cells 14 days after transduction with the indicated vectors. Mean fluorescence intensities (MFI) are indicated. **(B)** Validation of mCherry expression in 4-1BBL+ (blue) and 4-1BBL knockout (red) ACHN cells 20 days after lentiviral transduction with the hLUC-mCherry construct. Mean fluorescence intensities (MFI) are indicated. **(C)** Representative bioluminescence imaging of NXG immunodeficient mice bearing orthotopic ACHN tumors expressing or lacking 4-1BBL on day 27 post-graft. **(D)** Tumor growth over time, quantified by total flux (photons/second). Bioluminescence values were normalized to the signal measured at day 6 post-graft for each mouse, and then scaled by 10⁶ for visualization purposes. Left: individual mouse traces. Right: mean ± SEM for 4-1BBL⁺ (blue, n=6) and 4-1BBL KO (red, n=4) groups.

### The lack of 4-1BBL in the tumor results in improved tumor growth in CD34-humanized mice

To assess the impact of tumoral 4-1BBL on tumor control, the two cell lines were grafted under the renal capsule of CD34-reconstituted HuMice in three independent experiments, totaling 26 humanized mice reconstituted with 4 different donors (supplemental figure 3). Tumors failed to develop in some mice, including all mice from donor 3; these animals were therefore excluded from the analysis. The lack of 4-1BBL leads to an increase in tumor growth relative to controls in mice that originated from two different donors (Figure 3A-B). This is also observed in an independent experiment with donor 4 (supplemental figure 3). At the end of the experiments, the kidneys are visually bigger in mice receiving the 4-1BBL-deficient cell line (Figure 3C) and their weight are superior to controls in almost all mice (Figure 3D), extending the bioluminescence data. Altogether, the results show that deleting 4-1BBL improves tumor growth and that this effect is mediated by the human immune system of CD34-humanized mice.

**Figure 3.**
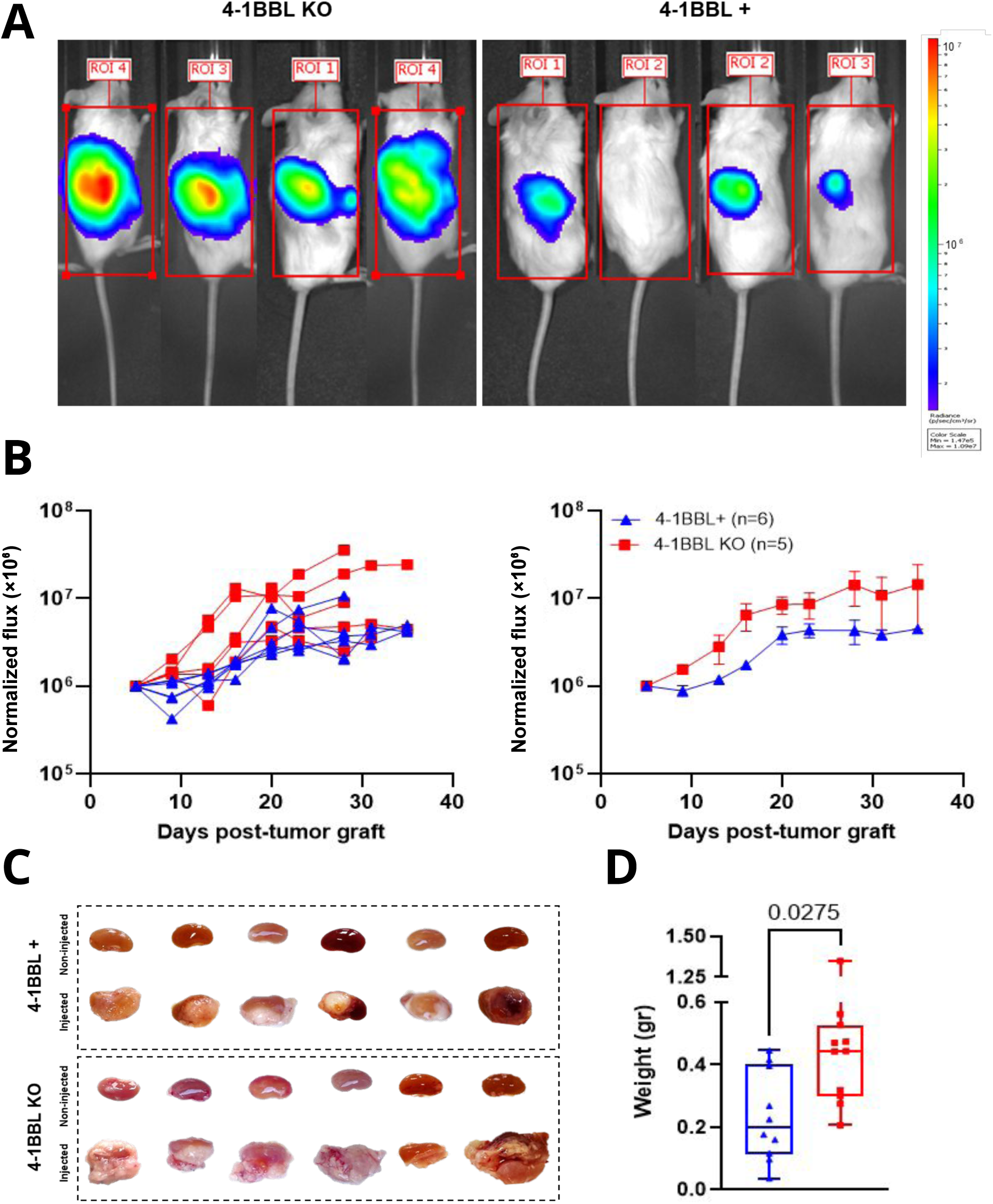
The lack of 4-1BBL in the tumor results in improved tumor growth in CD34-humanized mice. **(A)** Representative bioluminescence images of CD34 humanized NXG mice orthotopically engrafted with 4-1BBL-deficient (KO, left) or 4-1BBL-expressing (right) ACHN cells at day 25 post-graft **(B)** Quantification of tumor growth in mice receiving 4-1BBL KO tumors (red) or 4-1BBL+ cells (blue) quantified by total flux (photons/second). Bioluminescence values were normalized to the signal measured at day 6 post-graft for each mouse, and then scaled by 10⁶ for visualization purposes. Left: individual traces. Right: average signal ± SEM **(C)** Ex vivo analysis of tumor masses collected at day 35 post tumor-graft. Left: macroscopic images of tumors from 4-1BBL⁺ (top) and KO (bottom) groups. Right: tumor weights in the 4-1BBL KO (red) vs the 4-1BBL+ group (blue). Statistical significance was assessed using an unpaired *t*-test.

### Reduced immune aggregates in the tumor in the absence of 4-1BBL

To decipher the immune mechanism at play that may explain this improvement, we analyzed immune infiltrates by immunofluorescence staining of CD3, CD20, FOXP3 and CD8 (Figure 4). The analysis shows a marked presence of human immune cells within the tumor-bearing tissue, irrespective of the 4-1BBL status of the tumor, and contrasting with the scarce or negligible infiltration observed in non-tumor-bearing tissue (Figure 4A). We next determined the amount of immune aggregates using automated classification of immune cells density within a defined area (Figure 4B). A decrease in the surface occupied by clustered immune cells labeled with CD3, CD20, FOXP3 and/or CD8 over the total surface of the examined area is readily visible in the absence of 4-1BBL (Figure 4B). Overall, a decrease in the frequency of these immune aggregates per surface unit is observed at 4 different locations in the kidney (Figure 4C). However, the cellular content of these aggregates is the same irrespective of the 4-1BBL status of the tumor (Figure 4D). Thus, 4-1BBL deletion results in bigger tumors with a concomitant reduction of immune aggregates in the kidney but with no major recomposition of these aggregates.

**Figure 4.**
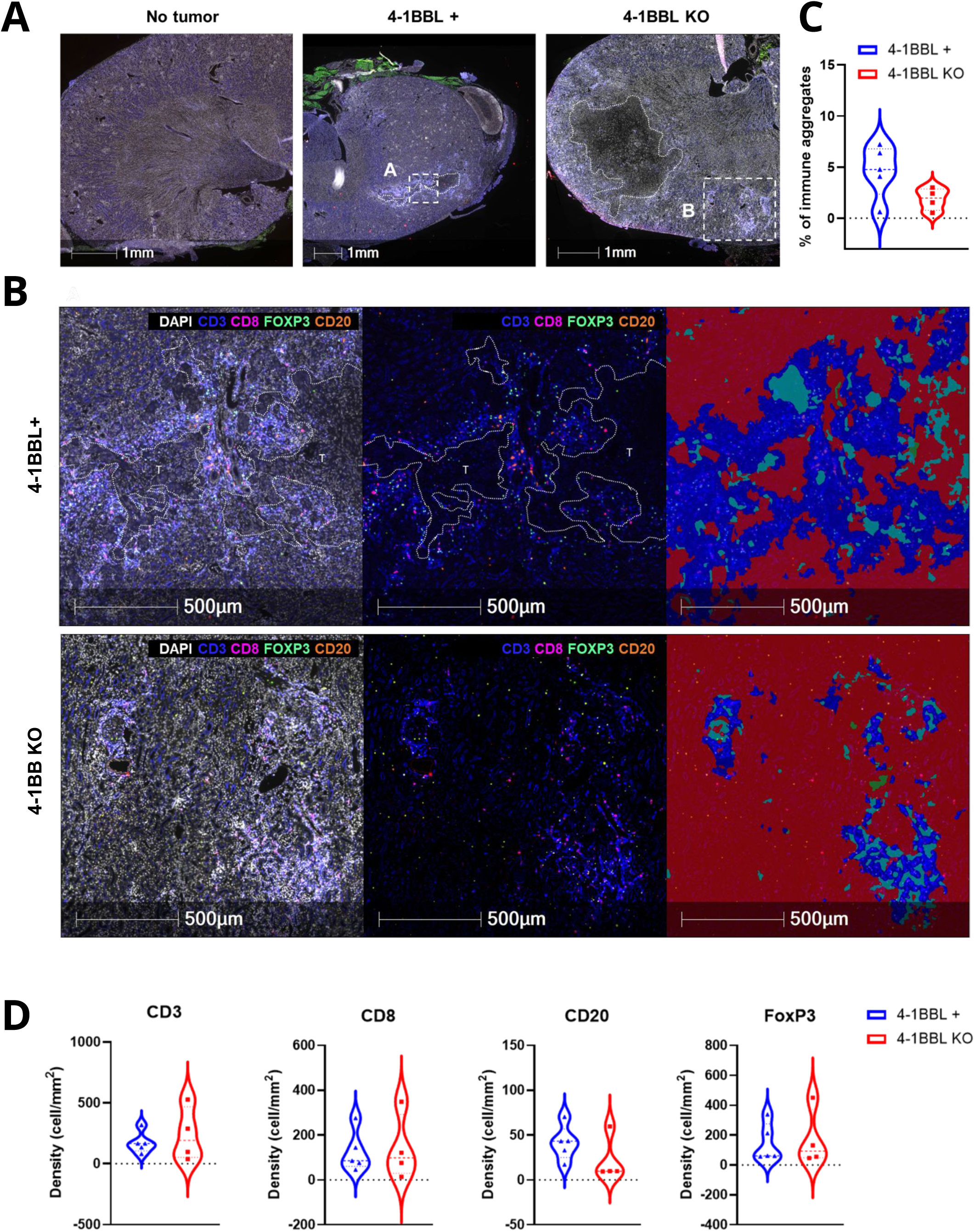
Reduced immune aggregates in the tumor in the absence of 4-1BBL. **(A)** Representative immunofluorescence images of kidney sections from mice with no tumor (left), implanted with 4-1BBL^+^ (middle) or with the 4-1BBL KO (right) ACHN tumor cells. Dotted areas indicate the regions used in (B). Scale bars of 1mm are shown. **(B)** Representative staining for CD3 (blue), CD8 (pink), FoxP3 (green), CD20 (orange) and DAPI (white) in regions outlined in (A). Tumor areas are outlined with dashed white lines and labeled “T”. Rightmost panels show HALO-AI based computational segmentation used to quantify immune aggregates : Empty or low cellularity regions appear in red and light blue while densely infiltrated areas regions appear in dark blue. Scale bars of 0.5mm are shown. **(C)** Quantification of immune aggregates (expressed as % of tissue areas) based on the HALO-AI classification in (B). Each dot represents an individual tissue section from the tumor-bearing kidney of one animal for each group. **(D)** Density of immune cell subsets within the immune aggregates, measured as number of cells per mm^2^ for CD3^+^, CD8^+^, CD20^+^ and FOXP3^+^ cells in 4-1BBL+ (blue) and 4-1BBL KO (red) tumors.

### The lack of 4-1BBL on the tumor impacts proliferation and exhaustion of 4-1BB+ CD8 T cells

Multiparameter flow cytometry was used to further dissect the tumor immune landscape, enabling the precise identification of multiple immune subsets (supplemental figure 4A). There is no significant difference in the proportions (supplemental figure 4B) nor numbers (supplemental figure 4C) of T, B, NK, or Treg cells within the tumor regardless of the 4-1BBL status. The immune composition of the draining lymph node and of the spleen is also identical regardless of the 4-1BBL status of the tumor (data not shown). In the tumor, the proportion of 4-1BB^+^ Tconv and Treg cells are decreased whereas the proportion of 4-1BB^+^ CD8 T cells is increased with the 4-1BBL KO tumor (Figure 5A-B). Other immune subsets expressing 4-1BB are not affected by the deletion (Figure 5B). Only 4-1BB^+^ CD8 T cells frequencies are increased in the tumor draining lymph nodes (dLN) but not in the spleen (supplementary figure 5A). Proliferation of 4-1BB^+^ cells in the tumor follows the same trends as 4-1BB expression, i.e decreased in Tconv and increased in CD8 T cells (Figure 5C). Importantly, proliferation of 4-1BB^-^ cells is not affected in the tumor, whatever the subset (supplemental figure 5B), indicating that proliferation of 4-1BB+ cells is driven by 4-1BBL expressed by other cells than the tumor. Similar to 4-1BB expression, proliferation of CD8 T cells is increased in the dLN but not in the spleen (supplemental figure 5C). Regarding 4-1BB^+^ Treg, their proliferation is not influenced by the tumor’s 4-1BBL status (Figure 5C), and their immunosuppressive phenotype remains unchanged (supplemental figure 5D). The proliferation of CD8 T cells in the absence of 4-1BBL coincides with an enrichment of PD-1⁺CD69⁻ cells expressing granzyme-B, indicative of a combined exhausted and cytolytic differentiation (Figure 5D).

**Figure 5.**
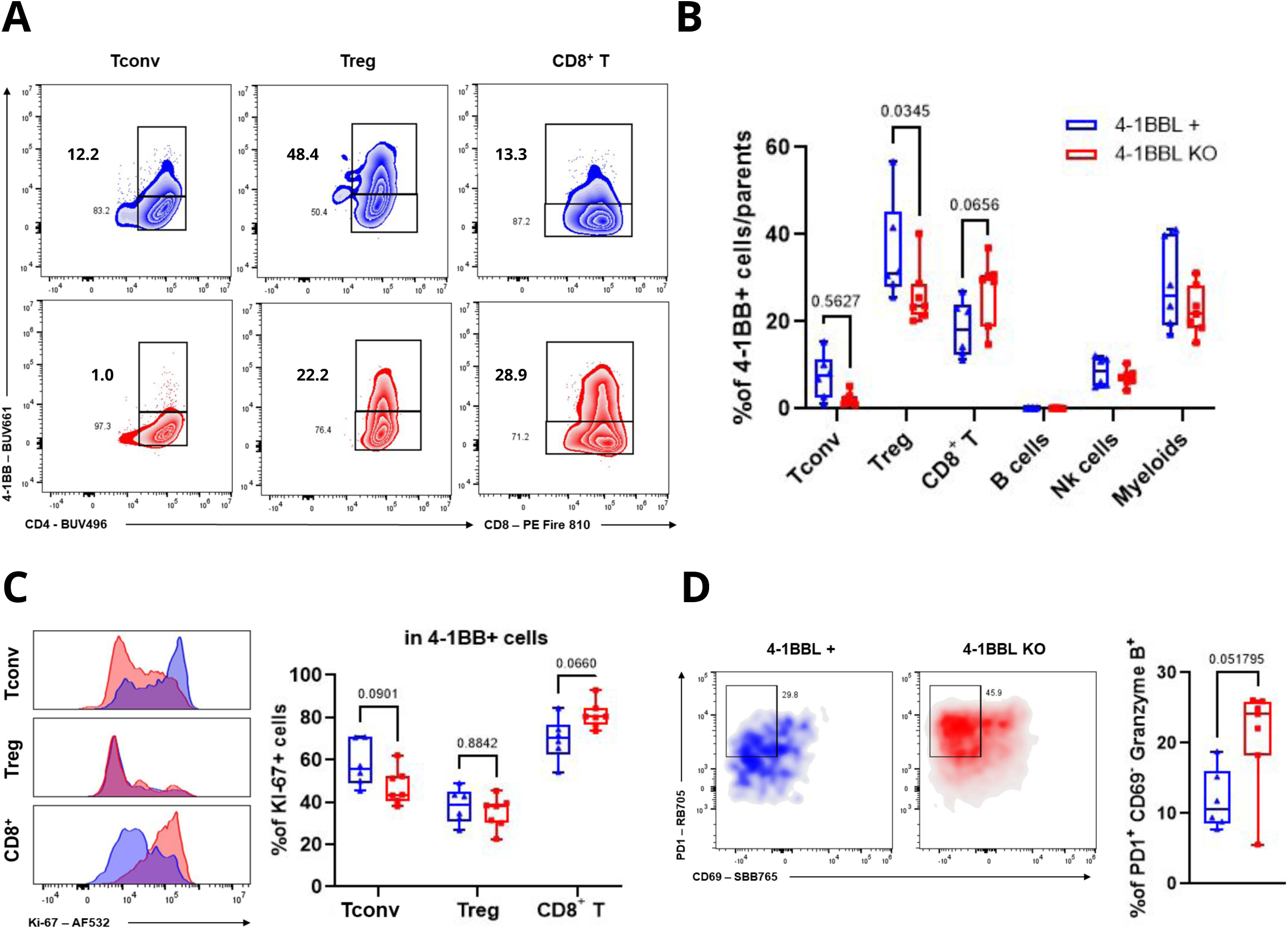
The lack of 4-1BBL on the tumor impacts proliferation and exhaustion of 4-1BB+ CD8 T cells. **(A)** Representative profiles of 4-1BB expression among CD4^+^FOXP3^-^CD25^-^ (Tconv), CD4^+^FOXP3^+^CD25^+^ (Treg) and CD8^+^ T cells isolated from 4-1BBL+ (blue) or 4-1BBL-deficient (red) tumors. The frequencies of cells in each quadrant are indicated **(B)** Frequencies of 4-1BB^+^ cells among the indicated immune subsets in the indicated group of mice in the tumor **(C)** Proliferation of 4-1BB^+^ cells in each T subsets in the tumor assessed by Ki-67 expression measurement. Left : representative histogram for Ki-67 in the indicated subset from 4-1BBL^+^ (blue) and 4-1BBL KO (red) tumors. Right : quantification of KI-67+ cells among each 4-1BB^+^ T cells subsets. (D) Co-expression of PD-1, CD69 and Granzyme B among CD8^+^ T cells in the tumor. Left : representative density plot of CD8^+^ T cells stained for PD1 and CD69 in 4-1BBL^+^ (blue) and 4-1BBL KO (red) tumors. Right : Frequency of CD8⁺PD1⁺CD69^-^Granzyme B⁺ cells among CD8^+^ T cells. Each dot represents an animal. Statistical testing of the null hypothesis was performed using one-way ANOVA on panel (B) and (C) and Mann-Whitney two-ways unpaired test for the panel (D).

### The lack of 4-1BBL in the tumor induces exhaustion of cytotoxic CD8 T cells

To confirm and extend this observation, we purified total human CD45^+^ cells from the tumor of three mice grafted either with the 4-1BBL+ or KO ACHN cell line and performed single cell RNA sequencing with a primer panel targeting close to 400 genes of the immune response (Figure 6). Seven distinct clusters emerge from the UMAP analysis of the data (supplementary figure 6A). The major immune subsets are readily identified by classical lineage markers: CD8A and CD8B for CD8 T cells, TRAT1, IL7R and CD5 for CD4+ T cells, FOXP3 and IL2RA for Treg (supplemental figure 6B). A cluster of myeloid cells defined by CD14 and CD33 is also detected. Moreover, two subsets of B cells differing on their putative ability to produce immunoglobulins are identified. Myeloid and B cells will not be further analyzed here.

**Figure 6.**
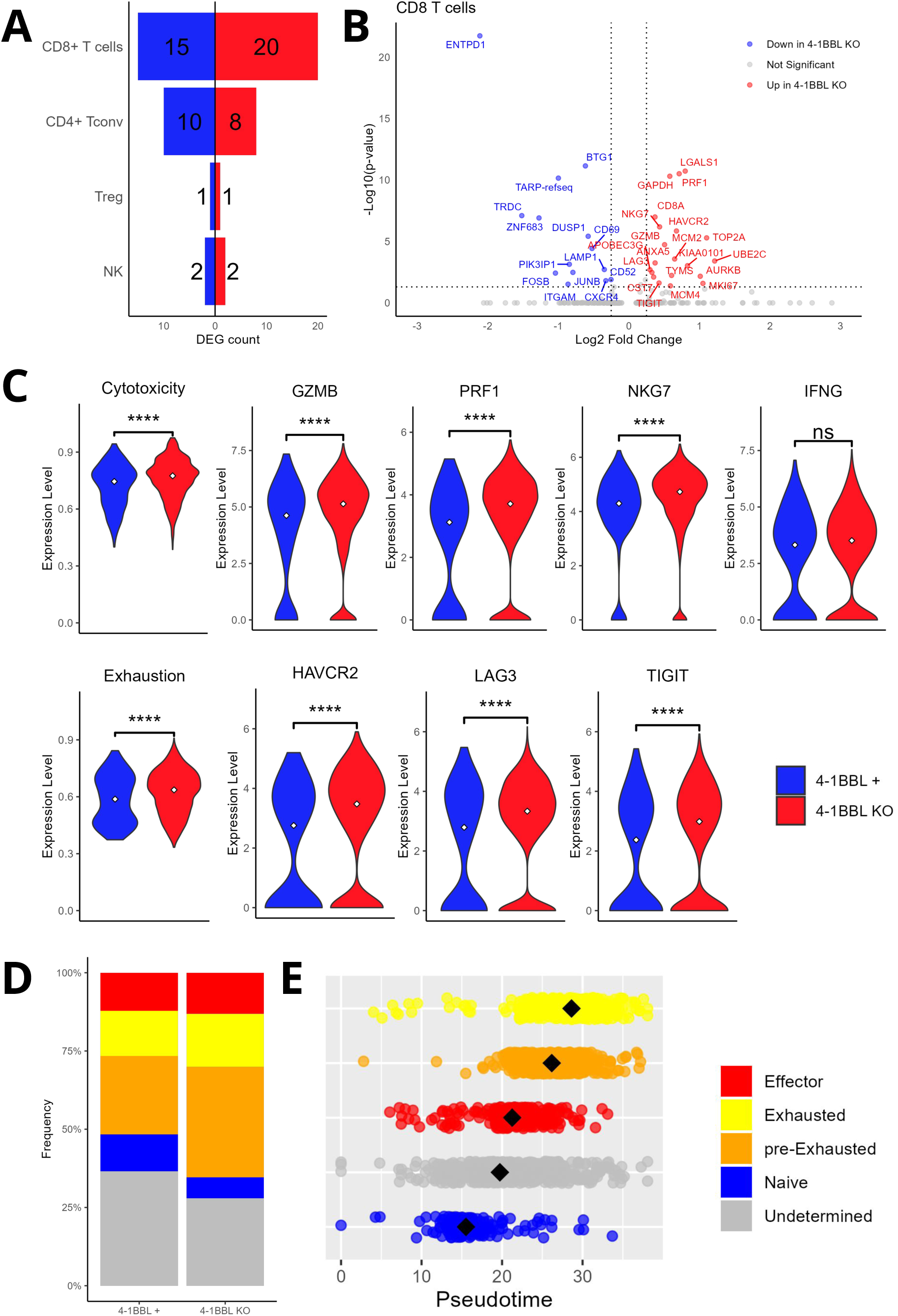
The lack of 4-1BBL in the tumor induces exhaustion of cytotoxic CD8 T cells. **(A)** Number of differentially expressed genes (DEG) significantly upregulated (red) or down-regulated (blue) in intratumoral CD8^+^ T cells, Tconv, Treg, and NK cells from 4-1BBL+ or 4-1BBL KO tumors. Each immune cell cluster was defined based on the annotation shown in supplementary figure S6. **(B)** Volcano plot showing DEG in CD8+ T cells from 4-1BBL+ versus 4-1BBL KO tumors. **(C)** Violin plots showing module scores for cytotoxicity and exhaustion gene signatures in CD8⁺ T cells alongside the expression levels of some genes for each signature. GZMB, PRF1, NKG7, and IFNG for the cytotoxicity signature and HAVCR2, LAG3, and TIGIT for the exhaustion signature. Medians are represented by a diamond shape. **(D)** Distribution of CD8⁺ T cells across distinct functional states defined by enrichment in gene signature : Effector (cytotoxic), Exhausted, pre-Exhausted (co-expression of cytotoxic and exhaustion signatures), Naive, and Undetermined (no signature enrichment) in CD8 T cells from 4-1BBL+ or 4-1BBL KO tumors. **(E)** Pseudotime trajectory illustrating the distribution of CD8⁺ T cell states defined in D. Each dot represents a cell. For panels A and B, the significant threshold is an adjusted p-value < 0.05 and a log2 fold change > 0.25 or < -0.25. A Wilcoxon test was used for panel C. ****: p-value < 0.0001.

Differential gene expression analysis within each of those clusters was performed according to the 4-1BBL status of the tumor. The analysis reveals limited changes in the transcriptome of lymphoid cells in the absence of 4-1BBL (Figure 6A). Most of those changes concern CD8 T cells whereas no significant changes are observed in the transcriptome of Treg and only moderate changes are observed for CD4 T cells, showing that the absence of 4-1BBL in the TME mostly affects the CD8 T cell transcriptome (Figure 6A). Detailed examination of the volcano plot for CD8 T cells reveals increased expression of TIM-3 (HAVRC2), LAG-3, TIGIT exhaustion markers along with enhanced expression of cytotoxicity-associated genes such as GZMB (Granzyme-B), PRF1 (Perforine) and NKG7 but not LAMP1 (Figure 6B). Thus, based on multiple reference gene signatures, we identified a significant alteration in the exhaustion and cytotoxic profiles of CD8 T cells in the absence of 4-1BBL (Figure 6C). This is confirmed at the level of the expression of single genes listed above. A noticeable exception is the untouched expression of IFNg in the absence of 4-1BBL in the TME (Figure 6C). Similar levels of IFNg production by CD8 T cells of the tumor is also observed at the protein level in the two groups of mice after in vitro PMA/Ionomycine restimulation (data not shown). Then, clusters of naïve, effector, pre-exhausted and exhausted CD8 T cells were defined based on the expression of gene signatures. The frequency of pre-exhausted CD8 T cells is increased in tumors lacking 4-1BBL (Figure 6D). As expected, this cluster appears as an intermediate between effector and fully-exhausted CD8 T cells in a pseudo time analysis with naïve cells as the reference (Figure 6D).

### CD8 T cells express high levels of 4-1BBL in the tumor and reduced AP1 signaling

So far, we have shown that the ACHN cell line deprived of 4-1BBL is associated with a surprising increase in the proliferation and proportion of CD8 T cells that only affects 4-1BB+ cells. This observation suggests that the lack of 4-1BBL co-stimulation by the tumor might be compensated by 4-1BBL co-stimulation by other cells. We therefore determined which cell type could be responsible for that compensation. Surprisingly, the highest level of 4-1BBL in the tumor is observed on CD8 T cells themselves with only scarce expression in myeloid or B cells, regardless of the 4-1BBL status of the tumor (Figure 7A). A slight increase in the proportion and in the intensity of 4-1BBL expression is observed on CD8 T cells in the absence of 4-1BBL in the tumor (Figure 7B). A similar observation is made from a published dataset of single cell RNA sequencing data (26): CD8 T cells and the tumor itself are major cell types expressing 4-1BBL at the mRNA level in ccRCC. Moreover, a discrete but well-defined cluster of 4-1BBL+ CD8 T cells is observed in a cluster of 4-1BBhi CD8 T cells (supplementary figure 7B), showing co-expression of 4-1BB and 4-1BBL. Thus, CD8 T cells expressing 4-1BBL in our CD34-humanized mouse mode are also observed in human kidney cancer patients, adding strength to the model. In the absence of 4-1BBL in the tumor, it is most likely that 4-1BBL expressed by CD8 T cells themselves is responsible for the 4-1BB-dependent effects on proliferation and exhaustion.

**Figure 7.**
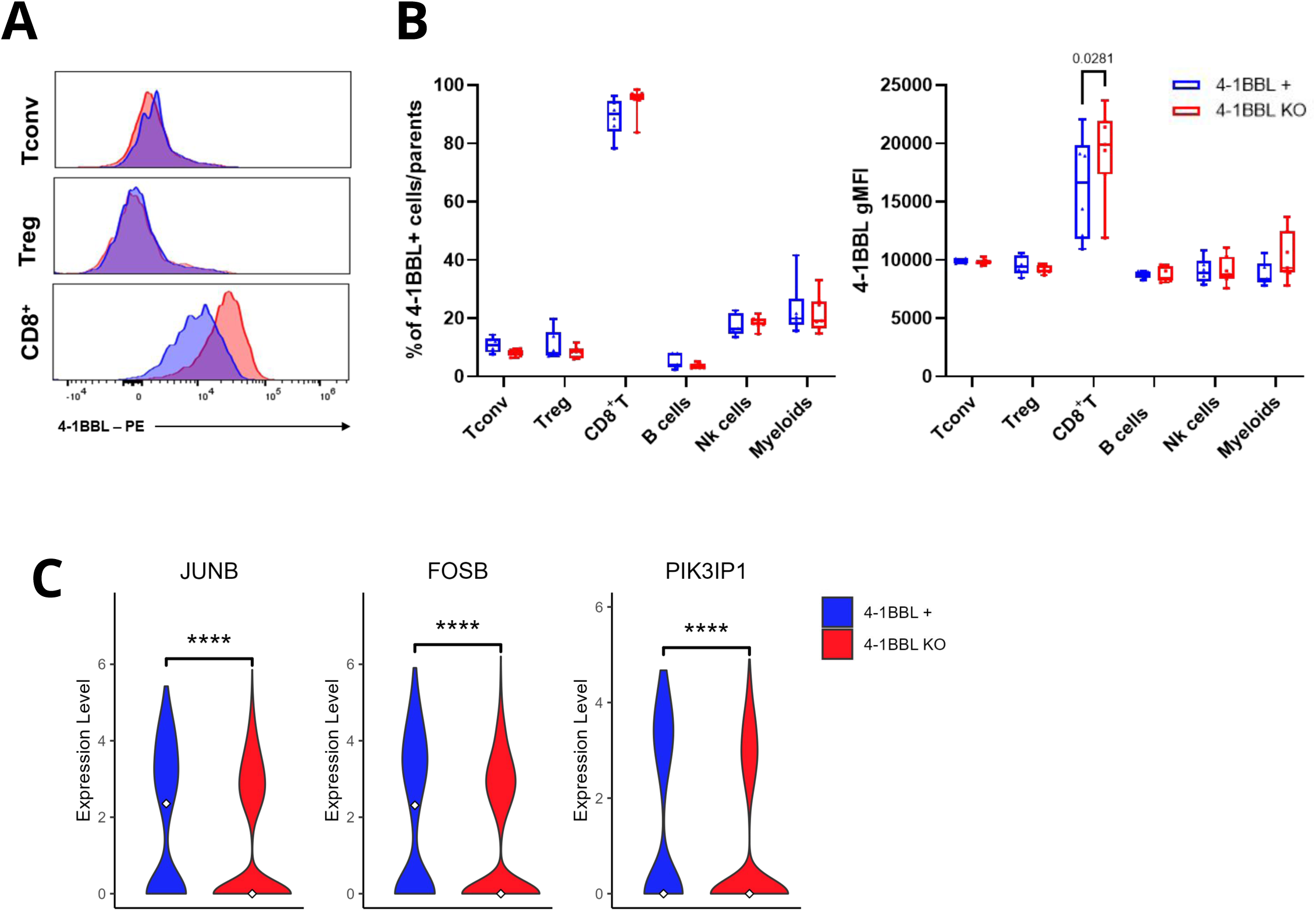
CD8 T cells expressed high levels of 4-1BBL in the tumor and reduced AP1 transcription factors. **(A)** Representative profile of 4-1BBL expression in the indicated subsets **(B)** Flow cytometry quantification of the frequencies of 4-1BBL+ cells (right) and the mean fluorescence intensity (MFI) (left) in the indicated subsets in tumors expressing (blue) or lacking 4-1BBL (red). (C) Violin plot showing differential expression of the transcription factors *JUNB*, *FOSB*, and the negative regulator *PIK3IP1* in CD8^+^ T cells from 4-1BBL+ (blue) or 4-1BBL KO (red) tumors. Statistical comparison in (B) were performed using the Wilcoxon-test; ****: p-value < 0.0001.

Finally, we investigated which signaling pathway could be associated with proliferation and exhaustion of CD8 T cells in our single cell RNA sequencing data. We observe lower expression of two AP-1 signaling members, FOSB and JUNB, and of the PI3K signaling pathway inhibitor, PIK3IP1 in the CD8 cluster (Figure 7C).

Overall, abolishing 4-1BBL co-stimulation of CD8 T cells *in trans* increases their acquisition of 4-1BB and 4-1BBL, improves their proliferation, and induces their differentiation into an exhausted/cytotoxic phenotype with reduced expression of AP1 transcription factors and an improved PI3K signaling pathway.

## Discussion

Previous attempts at modeling kidney cancer with Patient Derived Xenografts (PDX) lacked the immune component, limiting therapy research to chemotherapy with no applications to immunotherapy (27). Although CD34-humanized mice can also be grafted with PDX (28), this was established for RCC with PBMC-grafted mice with limited relevance to human physiology (29). More recently, a CD34-humanized mice model of ccRCC was used to validate a novel therapeutic strategy for partially HLA-matched CAR-T cells secreting anti-PD-L1 antibody, but how the model reproduced the histopathological features of patients was not reported (30). To our knowledge, we report the first model of pRCC in a CD34-humanized mice model, carrying a human tumor in the kidney and a complete human immune system. Histological analysis revealed structures typical of pRCC but also of a sarcomatoid type, previously observed in subcutaneous xenografts of ACHN (25). Indeed, the ACHN cell line has been assigned a pRCC genotype based on comparison with patients, but did not differentiate as such in vivo upon subcutaneous implantation (25). Here, we show that the orthotopic graft of ACHN might mimic more closely the papillary nature of the cell line although this remains to be formally determined by additional phenotypic characterisation, such as expression of CK7 or AMACR. Nevertheless, we took advantage of this model to investigate the functional consequences of 4-1BBL deletion on cancer cells, focusing on its impact on CD8⁺ T cells and Tregs.

Our working hypothesis was that tumor-expressed 4-1BBL could activate Treg through 4-1BB, thereby creating a favorable TME to the tumor in which case removal of this pro-Treg checkpoint should have led to a better tumor control. A dominant role of 4-1BBL inTreg activation and suppression would have explained the clinical data showing a correlation between 4-1BBL expression and poor survival of pRCC patients. Clearly, this hypothesis does not hold with the current model. This negative result does not rule out that Treg may contact the tumor directly but the few genes that are modified in Treg in the context of the 4-1BB KO cell line does not favor this hypothesis. Only 4-1BB+ Treg were affected by the deletion of 4-1BBL in the tumor. Thus, 4-1BBL expression by the tumor is either necessary to maintain 4-1BB expression by Treg or is necessary for 4-1BB+ Treg survival. Although abolition of 4-1BB signaling has been shown to affect Treg suppressive functions (13,16), 4-1BB can also promote their proliferation (31). The role of 4-1BB co-stimulation on Treg survival is less clear. Either way, the overall proportion of Treg and their immunosuppressive phenotype were not affected by the deletion, in contrast to tumor growth that was enhanced if expression of 4-1BBL by the tumor was lacking. Thus, expression of 4-1BBL by the tumor does not have a major impact on Treg activation and function in our model.

In contrast, our results strongly suggest that the lack of 4-1BBL on the tumor affects the effector arm of the immune response, and more precisely CD8 T cells. However, the improved tumor growth is not correlated to reduced cytotoxic CD8 T cells. On the contrary, we observe several quantitative and somewhat paradoxical phenotypic modifications due to the absence of 4-1BBL on tumor cells: increased proliferation and acquisition of 4-1BB and 4-1BBL, as well as increased expression of cytolytic and exhaustion markers. Despite this upregulation of cytolytic markers, these T cells were dysfunctional, not capable of eradicating the tumor, thereby explaining the increase in tumor size. This dysfunction might be attributed to increased exhaustion of CD8 T cells, as indicated by elevated expression of TIM3, LAG3 and TIGIT on the same cells. We also noted that the degranulation marker LAMP1/CD107a was not increased in the absence of 4-1BBL, providing an hypothesis to explain the lack of efficacy of cytotoxic molecules expressed by CD8 T cells. A defect in LAMP1 was implied in the inefficacy of NK cells to kill their targets (32) and that may also be the case for dysfunctional CD8 T cells. Moreover, both our transcriptomic and protein analyses indicate that IFN-γ production is not increased in the absence of the ligand, supporting the hypothesis that IFN-γ may be a key effector molecule for tumor control in our model.

What could be the mechanism that may explain this novel function of 4-1BBL? To our knowledge, we report for the first time that CD8 T cells are the main immune subset expressing 4-1BBL in the TME, at much higher levels than any other subsets, B cells and myeloid cells included. The expression of 4-1BBL on CD8 T cells is even higher in the absence of 4-1BBL on the tumor. In those conditions, a reduced AP-1 and an improved PI3K signaling pathway are observed, manifested by a reduction in the expression of FOSB and JUNB and PIK3IP1. Reduced AP-1 signaling pathway is often observed in exhausted T cells that might be due to binding of unbound NFAT:AP1 complex on co-inhibitory genes (33,34). Also, inhibition of the PI3K signaling pathway improves CD8 function and prevents exhaustion (35). Thus, the transcriptomic profile of AP-1 and PI3K signaling pathways observed in the absence of 4-1BBL in the TME is compatible with the engagement of CD8 T cells into functional exhaustion. Because the absence of 4-1BBL expression in the tumor is likely to reduce trans-signaling of 4-1BB in CD8 T cells, we conclude that *cis*-activation of 4-1BB by 4-1BBL may induce the transition of functional cytotoxicity into exhaustion that we describe here. In line with our hypothesis, a recent report describes improved proliferation and increased expression of cytotoxic markers in human CD8+ T cells stimulated with 4-1BB in *cis* compared to *trans* (21). The impact on functional exhaustion was not determined in this study. A recent study in mice using 4-1BB- and 4-1BBL-deficient cells indirectly showed that *cis*-interaction is necessary for CD8 T cell survival and to enhance the migration into the tumor dLN with a resulting effect on MC38 tumor growth (36). If our hypothesis is correct, limiting *cis*-activation of CD8 T cells with antibodies blocking 4-1BBL may limit T cell exhaustion and favor tumor rejection of the 4-1BBL-KO cell line. Our humanized mice model of kidney cancer will be invaluable to test this hypothesis.

Recently, it was proposed that 4-1BB signaling on CD8 T cells favored terminal differentiation and improved tumor control (12), a result that may seem at odds with our hypothesis. First, it should be stressed that the origin of 4-1BBL co-stimulation was not investigated in the work of Pichler et al. that mostly relies on 4-1BB agonists in syngeneic murine models. Here, we clearly identify 4-1BBL on the tumor cells as a major checkpoint with a novel biological function, that is preventing immune exhaustion of human CD8 T cells. Second, our results suggest that the indubitable role of 4-1BB co-stimulation on CD8 T cell activation might be delivered in *trans*, which is also the case for agonists. In contrast, *cis*-activation would lead to exhaustion, as suggested here and by others (21). In the complex world of T cell co-stimulation, the two mechanisms may superimpose with a positive or negative impact on tumor growth depending on the context.

Our results might explain the poor prognosis of 4-1BBL in pRCC: high levels of 4-1BBL expression in the TCGA dataset may signify the presence of exhausted cytotoxic CD8 T cells expressing 4-1BBL, as shown here. It would be those exhausted 4-1BBL+ CD8 T cells that would be of poor prognosis in RCC, as previously shown (37,38). We show here that 4-1BBL expression level is also associated with bad prognosis in pancreatic cancer and many other malignancies. It would be interesting to investigate further whether this poor prognosis is linked to the presence of 4-1BBL+ exhausted CD8 T cells in the TME. It is also conceivable that 4-1BBL might act as a surrogate marker of biologically aggressive tumors, in which its upregulation reflects a detrimental immune activation program. For instance, it has been shown in lung cancer that 4-1BBL might instruct T cells to produce IFNg that in return increase the expression of PD-L1 by tumor cells (39).

Our results highlight the complexity of the 4-1BB/4-1BBL axis in solid tumors and underscore the need for further investigation into its functional role, cellular sources, and spatial dynamics in the renal tumor microenvironment.

## Materials and Methods

### Humanized Mice

NOD-Prkdc^scid^ -IL2^rgTm1^/Rj (NXG) mice, reconstituted with human CD34^+^ hematopoietic stem cells, were obtained at ∼20 weeks of age from Janvier Labs (Le Genest-Saint-Isle, France). Only female mice were used for all procedures. They were housed under specific-pathogen-free conditions in individually ventilated cages, with a 12-hour light/dark cycle at 22 ± 2 °C and 50 ± 10 % humidity. Food and water were provided ad libitum.

### Cell culture, generation and validation

ACHN cells were obtained from Cell Lines Services (Eppelheim, Germany) and cultured in DMEM (Gibco, Grand Island, NY, USA) supplemented with 10% fetal bovine serum, penicillin (100 U/mL) and streptomycin (100 µg/mL) at 37 °C, 5% CO₂.Mycoplasma contamination was excluded using the MycoAlert kit (Lonza, Basel, Switzerland). An ACHN hTNFSF9-KO (4-1BBL KO) clone was obtained from VectorBuilder (Guangzhou, China), generated by CRISPR/Cas9 RNP targeting exon 1 of TNFSF9; biallelic disruption was confirmed by PCR and Sanger sequencing. The ACHN 4-1BBL KO cells were seeded at 1.5 × 10^5 cells per well in 12-well plates containing 2 mL complete DMEM (Gibco, Grand Island, NY, USA) and incubated overnight. To reconstitute 4-1BBL expression, cells were transduced with the Lv-pCMV6-TNFSF9-GFP (OriGene, Rockville, MD, USA) lentiviral vector with a multiplicity of infection (MOI) of 10 (10 viral particles per cell) in 1 mL medium supplemented with 8 µg/mL polybrene (Sigma-Aldrich, St. Louis, MO, USA). Four hours post-infection, medium was topped up and cells were incubated for 16 h before exchange. Forty-eight hours after infection, GFP⁺ cells were isolated by FACS (BD FACSAria II; BD Biosciences) using GFP and forward/side scatter gates to achieve > 95% purity. An empty-vector (GFP-only) transduction served as a control.

The resulting 4-1BBL⁺ and 4-1BBL KO GFP⁺ clones were next transduced with LPP-hLUC-Lv214-025-C (OriGene Technologies) at MOI 10 under identical conditions to generate the final ACHN 4-1BBL⁺ GFP⁺ mCherry⁺ luciferase⁺ line. Stable expression of 4-1BBL and luciferase activity in both engineered and control lines was confirmed by flow cytometric detection of 4-1BBL and in vitro bioluminescence assays, respectively. Post-sort, all cell lines were maintained in complete DMEM without antibiotic selection for downstream experiments.

### Renal Subcapsular Implantation and tumor monitoring

Tumor implantation was performed at approximately 20 weeks post-CD34⁺ transplantation, a time point chosen to ensure stable human hematopoietic reconstitution. Two weeks before implantation, peripheral blood was collected with puncture of the maxillary vein. Human reconstitution was determined by flow cytometry. Animals were then stratified into experimental cohorts with matched levels of human chimerism (data not shown). One hour before surgery, mice received meloxicam (5 mg/kg SC). On the day of surgery, mice were anesthetized via intraperitoneal injection of ketamine (100 mg/kg) and xylazine (10 mg/kg), and the left kidney was exteriorized via a small dorsolateral incision. A total of 2 × 10^6 ACHN-luciferase cells (either 4-1BBL⁺ or 4-1BBL KO), suspended in 20 µL PBS, were gently injected under the renal capsule using a 29-gauge insulin syringe. The kidney was returned to the peritoneal cavity, and the incision was closed in two layers (6-0 Vicryl for muscle, 5-0 poly for skin). Postoperative analgesia (meloxicam, 5 mg/kg SC) was administered at 24 h and 48 h postoperatively. Tumor growth was monitored by weekly bioluminescence imaging for 30 days post-implantation: mice received an IP injection of D-luciferin (150 mg/kg), were anesthetized with isoflurane, and imaged 10 min post-injection on an IVIS Spectrum system. The bioluminescent signal over the graft site was quantified using Living Image 4.8.2 (PerkinElmer, Hopkinton, MA, USA). At study end, mice were euthanized by intraperitoneal overdose of a ketamine (200 mg/kg) and xylazine (20 mg/kg) mixture, followed by exsanguination to permit harvest of organs for downstream analyses.

### Preparation of single**-**cell suspensions

At study end, mice were anesthetized by intraperitoneal overdose of ketamine (200 mg/kg) and xylazine (20 mg/kg). Rightafter, animals were perfused via the left ventricle with ice-cold PBS to remove circulating leukocytes to permit organ harvest for clean downstream analyses. Kidneys were excised and cut into ∼0.5-cm fragments, then transferred into 5-mL tubes containing the Tumor Dissociation Kit, Mouse (Miltenyi Biotec, Bergisch Gladbach, Germany) enzymes reconstituted in RPMI-1640 as per the manufacturer’s recommendations. Samples were incubated at 37 °C on an orbital shaker set to 120 rpm for 45 minutes. Following digestion, suspensions were passed through a 70-µm cell strainer, washed with PBS-3% FCS (300 × g, 7 min, 4 °C), and pelleted. Renal leukocytes were isolated by density centrifugation: cell suspensions were layered onto a 40%/80% Percoll gradient and centrifuged at 2,000 rpm for 20 minutes at 20 °C; mononuclear cells were collected at the interphase and washed twice in PBS-3% FCS. Splenic tissues and lymph nodes were mechanically disrupted through a 70-µm cell strainer to generate single-cell suspensions. For the spleen, residual erythrocytes were lysed by a 1-minute incubation in ammonium-chloride-potassium (ACK) lysis buffer, followed by washing in PBS-3% FCS. All cell suspensions were maintained on ice and counted prior to downstream applications.

### Hematoxylin and Eosin staining

Tissues were fixed in 4% paraformaldehyde (Electron Microscopy Sciences, Hatfield, PA, USA) at room temperature for 12 h, followed by dehydration and paraffin embedding. Five-micrometre sections of kidney were cut, deparaffinized, and stained with hematoxylin and eosin for 5 min at room temperature, then mounted and examined by bright-field microscopy (20X).

### Antibodies and flow cytometry

Single-cell suspensions in 96-well plates were incubated for 10 minutes at 4 °C with Human FcR Blocking Reagent (Miltenyi Biotec, Bergisch Gladbach, Germany), murine Fc block (2.4G2 supernatant) and LIVE/DEAD™ Blue fixable dye (Thermo Fisher Scientific, Waltham, MA, USA) to simultaneously assess viability and minimize non-specific binding. Surface markers were stained for 20 minutes at 4 °C in PBS-3% FCS using fluorochrome-conjugated antibodies listed in supplemental Table 1. For intracellular transcription factor detection, cells were fixed and permeabilized using the Foxp3/Transcription Factor Staining Buffer Set (eBioscience, San Diego, CA, USA), then stained for 45 minutes at 4 °C. Cytokine production was evaluated following a 4-hour stimulation with PMA (50 ng/mL) and ionomycin (5 µg/mL) in complete RPMI-1640 (10% FCS, 2 mM L-glutamine, 20 mM HEPES, 1 mM sodium pyruvate, 1 mM non-essential amino acids, 100 U/mL penicillin, 100 µg/mL streptomycin) supplemented with GolgiPlug (BD Biosciences, Franklin Lakes, NJ, USA). Data were acquired on a Cytek Aurora spectral cytometer and analyzed using FlowJo v10 (BD Biosciences, Franklin Lakes, NJ, USA). Spectral unmixing was performed using reference controls for each fluorochromes, allowing accurate deconvolution of overlapping emission spectra. The gating strategy used in the flow cytometry analysis is depicted in Supplementary Fig. 5.

### Single**-**cell capture, library preparation, and sequencing

Cell viability for all tumor tissue samples was above 90%, as assessed by flow cytometry using Calcein (BD Biosciences, USA) and DRAQ7 (Thermo Fisher Scientific, USA). Following the manufacturer’s instructions, six samples (three 4-1BBL^+^ tumors and three 4-1BBL KO tumors) were barcoded using BD Sample Tags and pooled for single-cell capture with the BD Rhapsody Single-Cell Analysis System (BD Biosciences, USA). Single-cell suspensions were loaded onto the BD Rhapsody Express system (BD Biosciences) and processed using the BD Rhapsody cDNA (cat. 633663) and Targeted Amplification (cat. 633664) kits per manufacturer’s instructions. A total of 20 000 cells were included in the pool. Briefly, captured mRNA was reverse-transcribed and amplified for 11 cycles using the pre-designed Immune Response Panel HS (299 primer pairs; cat. 633750). PCR1 products were purified with AMPure XP beads (Beckman Coulter, cat. A63881) and size-selected (350 to 800 pb for mRNA; ∼170pb for Sample Tag). Each fraction then underwent a second amplification (PCR2, 10 cycles). Resulting libraries were quantified by Qubit dsDNA HS assay (Thermo Fisher Scientific, Waltham, MA, USA; cat. Q32854) and assessed for fragment size on an Agilent 2100 Bioanalyzer (High Sensitivity DNA Kit; Agilent Technologies, Santa Clara, CA, USA; cat. 5067-4626). Libraries with concentrations > 1.5 ng/µL and expected size distributions (∼300–600 bp for mRNA amplicons) were normalized to 2.3 ng/µL (mRNA) or 0.1 ng/µL (Sample Tag), indexed by a final six-cycle PCR with Illumina primers, adjusted to 4 nM, pooled at a 93:7 ratio (targeted mRNA:Sample Tag), spiked with 15 % PhiX control, denatured, and sequenced (75 bp paired-end) on an Illumina NextSeq 2000 system with P1 reagents (100 cycles), as recommended by the manufacturer.

### Single-cell RNA Sequencing data processing and analysis

FastQ files were processed using the standard Rhapsody analysis pipeline (BD Biosciences) on Seven Bridges (sevenbridges.com), following the manufacturer’s recommendations. The resulting Seurat object was analyzed in R, and gene expression was log-normalized and read back into the original Seurat object. The normalized data were scaled in Seurat and principal components were calculated based on all genes present in the dataset. A JackStraw plot was then used to determine the significant principal components. The cells were then split into unsupervised clusters using a shared nearest neighbor modularity optimization-based clustering algorithm. Differentially expressed genes (DEG) for each cluster were identified using the Wilcoxon test in Seurat. Top DEG (filtered at the p < 0.05 and log₂ fold change >0.25 thresholds) were then used to identify different cell types. Similarly, DEG for CD8 T cells from 4-1BBL KO vs. WT were determined and plotted on a volcano plot. Signature enrichment analysis based on a naive signature (CCR7, LEF1, TCF7, SELL, IL7R, CD27, CD28, LTB and S100A10), a cytotoxic signature (GZMB, PRF1, NKG7, GZMA, GZMH, GNLY, IFNG and CX3CR1) and an exhausted signature (LAG3, PDCD1, CTLA4, TIGIT, HAVCR2, CXCL13, CXCR6 and IRF4), was performed using the Ucell R package, and pseudotime analysis was performed using the Slingshot R package. All data plotting was done with the ggplot2 R package.

### Multiplex immunofluorescence

Sections were deparaffinized with Clearene (Leica, catalog #3803600E), rehydrated in successive ethanol baths (100, 90, 70, 50 and then 0%), and subjected to heat-induced epitope retrieval with the target retrieval solution at pH 6.0 (Dako, catalog #S236784-2) or Tris–EDTA buffer (pH 8.0) for 30 min at 97 °C in a water bath. After cooling and two washes in TBS (Euromedex, catalog #ET220-B), endogenous peroxidases were quenched with 3% H₂O₂ for 30 min at room temperature, followed by a 30 min block in protein blocking solution (Agilent, catalog #X090930-2). For FoxP3 detection, anti-FoxP3 (clone 206D, mouse IgG1, Thermofisher, catalog #320102) was applied overnight at 4°C, washed in TBS-0.04% Tween 20, and followed by goat anti-mouse IgG1–HRP (Jackson Immunoresearch, catalog #115-035-205) for 30 min at room temperature. Tyramide-AF594 (ThermoFisher, catalog # B40957) was added as an amplification system after washing the tissue sections with TBS-0.04% Tween-20 and was incubated at room temperature for 10 min. The tissue sections were then incubated at 97°C for 10 min with the target retrieval solution after washing. The procedure was repeated several times to label the same tissue sections with various antibodies using tyramide conjugated to distinct fluorochromes for: anti-CD8 (clone SP16, rabbit polyclonal IgG, Thermofisher, catalog #MA5-14548) revealed with donkey F(ab’)2 anti-rabbit IgG-HRP (Jackson ImmunoResearch, catalog #711-036-152) and tyramide-AF647 (Thermofisher, catalog #B40958); anti-CD3 (rabbit polyclonal IgG, Dako, catalog #A0452) revealed with donkey F(ab’)2 anti-rabbit IgG-HRP (Jackson ImmunoResearch, catalog #711-036-152) and tyramide-CF430 (Interchim, catalog #96053); and anti-CD20 (clone L26, mouse IgG2a, Dako, catalog #M0755) revealed with goat anti-mouse IgG2a-HRP (Jackson ImmunoResearch, catalog #115-035-206) and tyramide-CF514 (Interchim, catalog # 92199). Nuclei were then stained with DAPI (1 µg/mL) (ThermoFisher, catalog # 62248) at RT for 5 min before coverslips were mounted onto glass slides with fluorescent mounting medium (Dako, catalog # S302380-2). All epifluorescent images were acquired on a Zeiss Axio Z1 fluorescent microscope (Carl Zeiss, Germany) using the Zen software. For mouse renal tumors, whole slide images were generated by scanning slides with a Zeiss Axio Z1 fluorescent microscope with adapted filters for imaging at LED 385nm, EmBP 450/40 for DAPI; 430nm, EmBP 480/40 for CF430; 511nm, EmBP 605/70 for CF514; 590nm, and EmBP 647/57 for AF594; 630nm, EmBP 690/50 for AF647. Scanning was performed at a magnification of 10x. Analysis was performed using Halo® software. After classifying the tumor regions, we performed nuclei segmentation and phenotypic analysis.

### Statistical Analysis

All statistical analyses were performed in GraphPad Prism v.10 (GraphPad Software, San Diego, CA, USA). The nature of the statistical test used to compare results is indicated in each figure legend. When necessary, the p-values of these tests are indicated in the figure panels. The statistical power of the analyses (alpha) was set arbitrarily at 0.05.

## Acknowledgements

The authors would like to thank Olivier Brégerie and Doriane Forest for taking care of the mice, Dr Nicolas Aubert and Etienne Camenen for help with bioinformatics, Aness Haddouche, Clara Cretet-Rodeschini for help with single cell RNA sequencing, Armanda Casrouge, and Solène Fastenackels for technical help and Pr Makoto Miyara, and Dr Jean-Luc Teillaud for support.

## Data availability statement

Data and source codes are available on request.

## Ethical statement

All experimental procedures were approved by the institutional animal care and use committee (APAFIS #40751-2023020116283147) and complied with the latest European guidelines. This study is reported in accordance with the applicable ARRIVE guidelines.

## Supplementary figure legends

**Supplemental figure 1. TNFSF9 is specifically overexpressed in kidney cancer. (A)** TNFSF9 (encoding 4-1BBL) shows cancer-enhanced expression, with highest RNA levels in kidney-derived cell lines presented as normalized transcripts per million (nTPM). **(B)** TNFSF9 mRNA expression is significantly elevated in renal tumors compared to other TCGA tumor types (UALCAN, TCGA). (source A-B: Human Protein Atlas (proteinatlas.org) (40) **(C)** Boxplot comparison of TNFSF9 expression between normal (gray) and tumor tissues (red) in three renal cancer subtypes: KICH (chromophobe), KIRC (clear cell), and KIRP (papillary) (UALCAN). * indicates statistical significance (p<0.05). The image was generated using XenaBrowser (xenabrowser.net) (41).

**Supplemental figure 2. Prognostic value of TNFSF9 expression across cancers. (A)** Upper panel: Kaplan–Meier overall survival analysis in patients with papillary renal cell carcinoma (KIRP) according to TNFSF9 expression. Lower panel: same analysis for clear cell renal cell carcinomas patients (KIRC). Data were filtered on the upper and lower 30% of all patients. Hazard ratio and p value are reported on the figure. Image was generated with TIMER 3.0 (compbio.cn/timer3) (42) **(B)** TNFSF9 expression increases with tumor stage progression, peaking at stage IV for KIRP (upper panel) and KIRC patients (lower panel) (one-way ANOVA analysis results are indicated) **(C)** Pan-cancer analysis of TNFSF9 prognostic impact using log-rank test. Bars represent significance levels (-log_10_ P values); red indicates longer survival with high TNFSF9, dark blue indicates shorter survival, and light blue indicates no significance. (source B-C: the Tumor Immune System Interaction database (cis.hku.hk/TISIDB) (43))

**Supplemental figure 3. Inter-individual variability in CD34-humanized mice.** ACHN expressing 4-1BBL (blue) or not (red) were grafted under the kidney capsule of CD34-humanized NXG mice reconstituted with 4 different donors. Tumor growth was quantified by bioluminescence (total flux, photons/second). Data were normalized to the baseline value measured at the initial time point.

**Supplemental figure 4. Spectral flow cytometry analysis of the tumors in CD34-humanized mice. (A)** Representative stainings and gating strategy to define lymphoid immune cells in the tumor of an ACHN-grafted humanized mouse. **(B)** Frequencies and **(C)** numbers per gram of tumor tissue of the indicated subset in 4-1BBL+ (blue) or 4-1BBL KO (red) in CD34-humanized mice 35 days after the grafts.

**Supplemental figure 5. Impact of 4-1BBL deletion in secondary lymphoid organs. (A)** Proliferation of 4-1BB-negative cells in each T subsets in the tumor assessed by Ki-67 expression measurement. **(B)** Frequencies of 4-1BB+ cells in the indicated immune subsets in tumor draining para-aortic lymph node (dLN) and in the spleen of CD34-humanized mice 35 days after the graft of 4-1BBL+ (blue) or 4-1BBL KO (red) ACHN cells. **(C)** Proliferation of CD8⁺ T cells measured by Ki-67 expression in dLN and spleen. **(D)** Expression of immune regulatory markers among FOXP3⁺CD25⁺ Treg in 4-1BBL+ (blue) or 4-1BBL KO (red) ACHN tumors. Each dot represents an individual mouse. Statistical comparisons were performed using one-way ANOVA.

**Supplemental figure 6. Cell type annotation from the single-cell RNA sequencing. (A)** UMAP of human CD45^+^ cells sorted from 4-1BBL + and 4-1BBL KO tumors in CD34-humanized mice 35 days after the grafts. Colors represent unbiased cell classification via graph-based clustering. Each dot represents an individual cell. (B) Heatmap for the top 10 genes with the highest expression enrichment in each cluster.

**Supplemental figure 7. CD8 T cells are the major cell type expressing 4-1BBL in clear cell renal cell carcinoma. (A)** Left panel: UMAP projection of single-cell RNA-Seq data from human tumor samples of ccRCC with cells colored by major lineage identity identified in the legend (retrieved from (26)). Right panel: TNFSF9 transcript abundance is indicated on a color scale (log-normalized expression). Arrows point to 4-1BBL-expressing cells in clusters of tumor cells and of lymphoid cells **(B)** Distribution of 4-1BBhi and 4-1BBlo CD8 T cells in the lymphoid cluster. Images were generated on the Single Cell portal of the Broad Institute (singlecell.broadinstitute.org) (44).

