## Supplementary figures and images for "The tumor necrosis superfamily member 4-1BBL expressed by tumor cells prevents exhaustion of CD8 T cells in a humanized mouse model of papillary renal cell carcinoma"

### supplemental figures

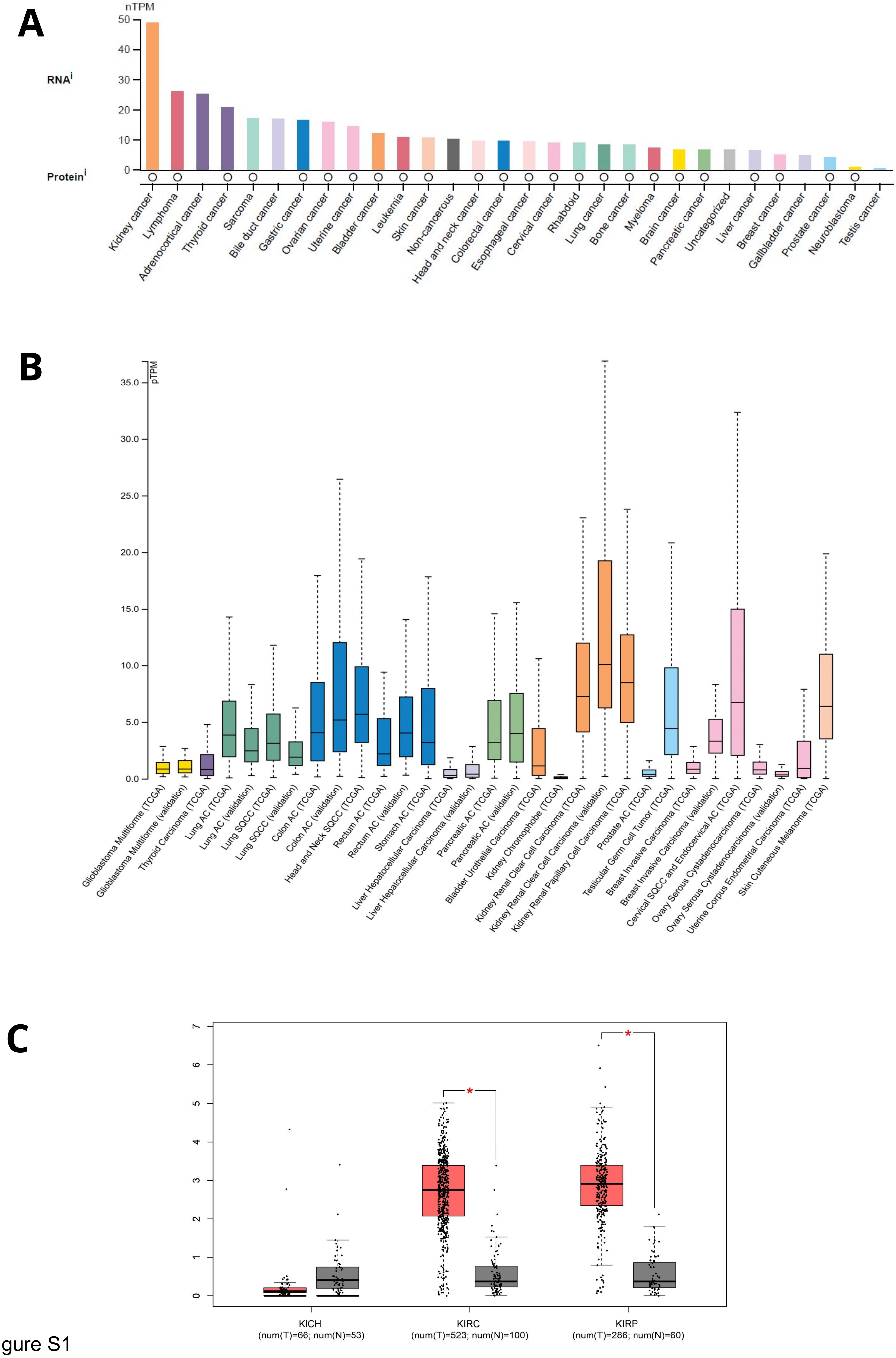

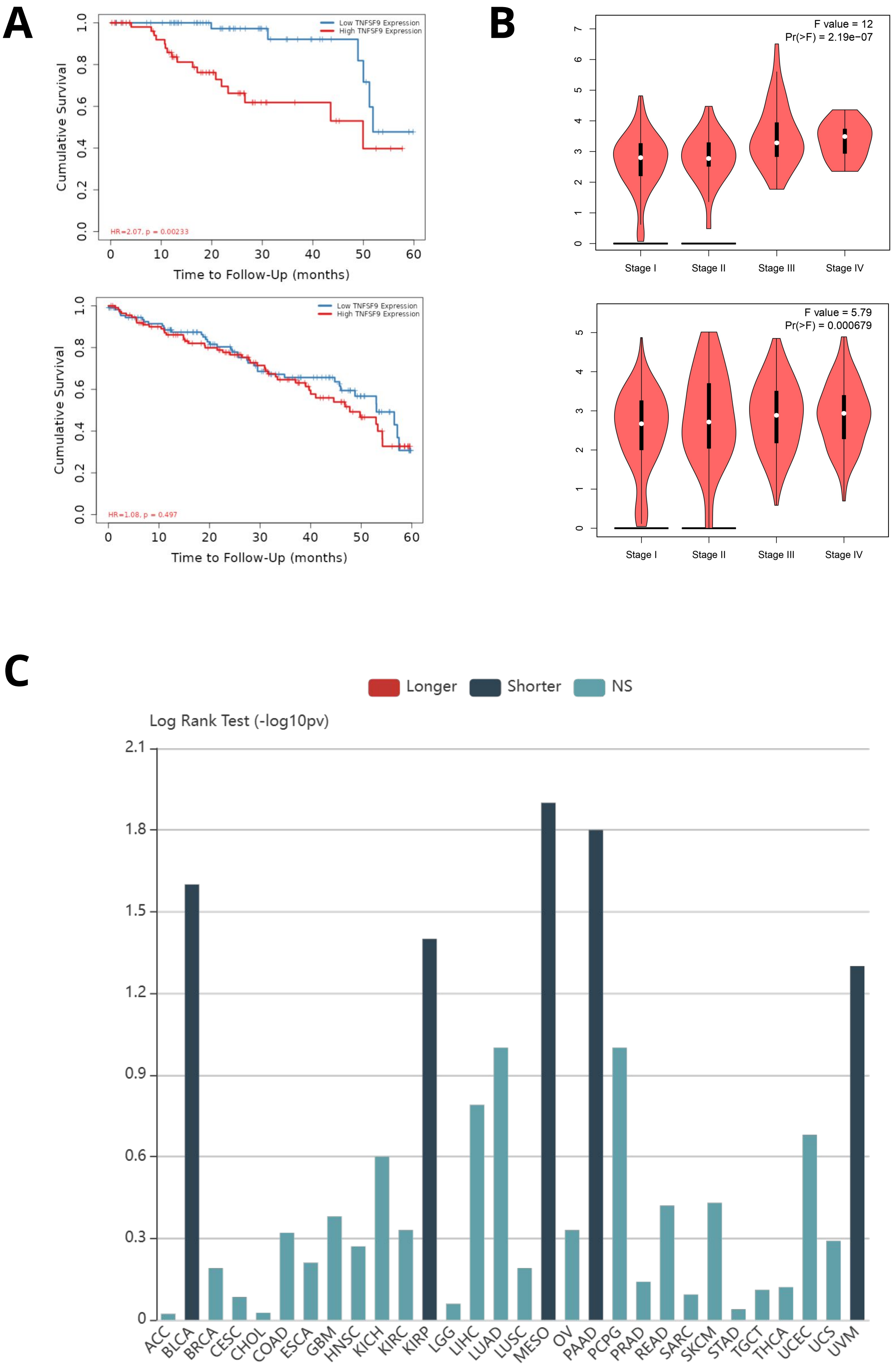

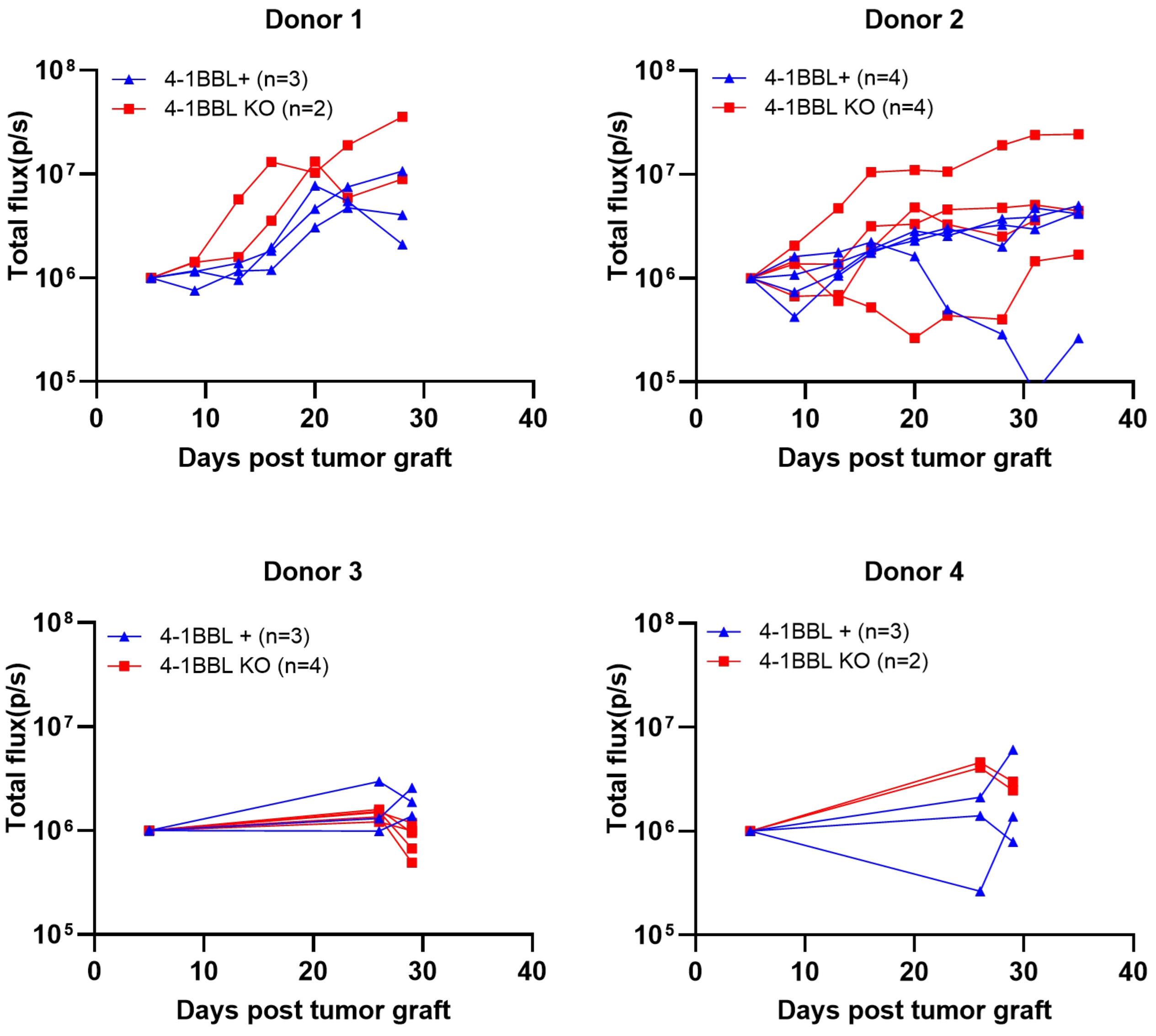

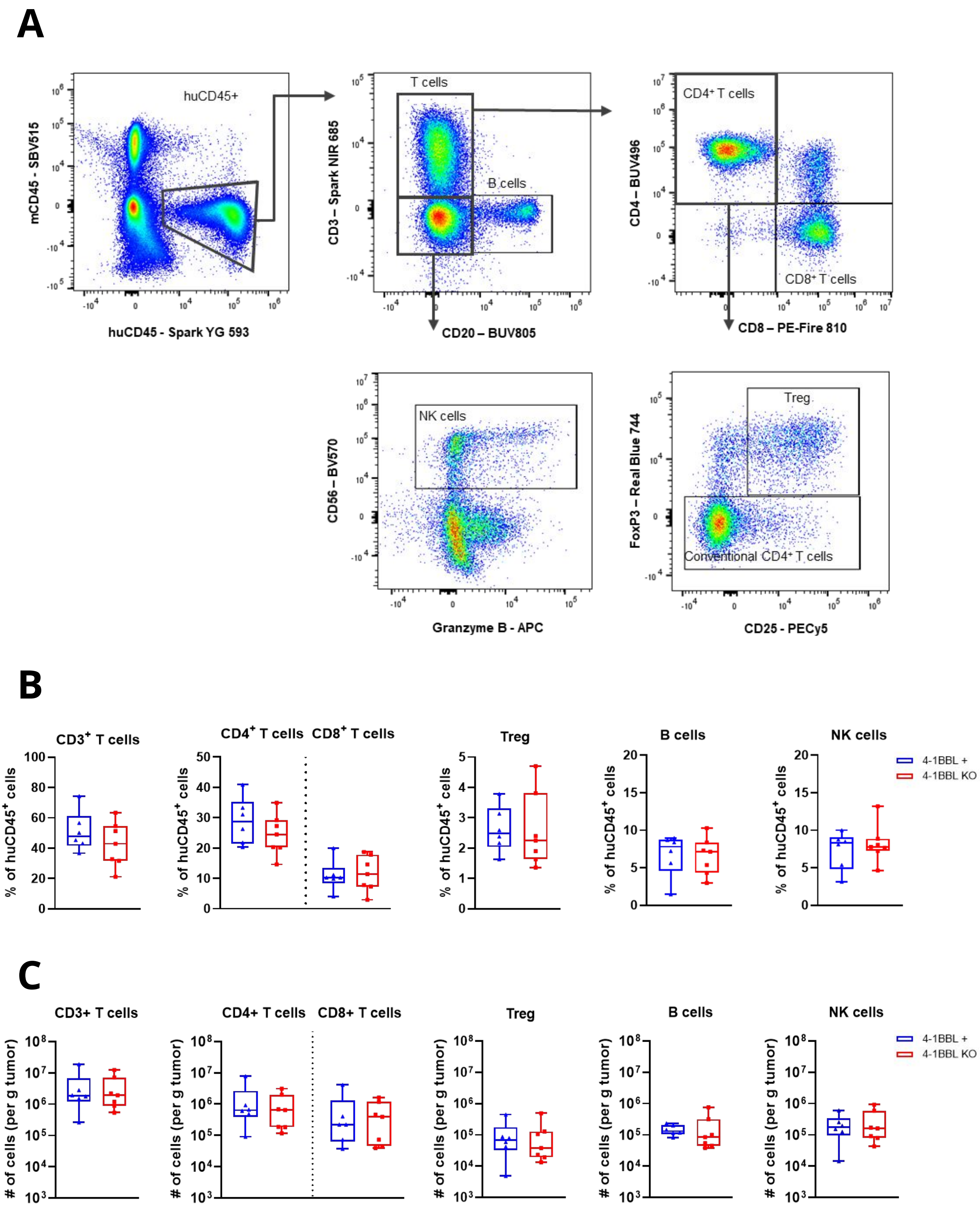

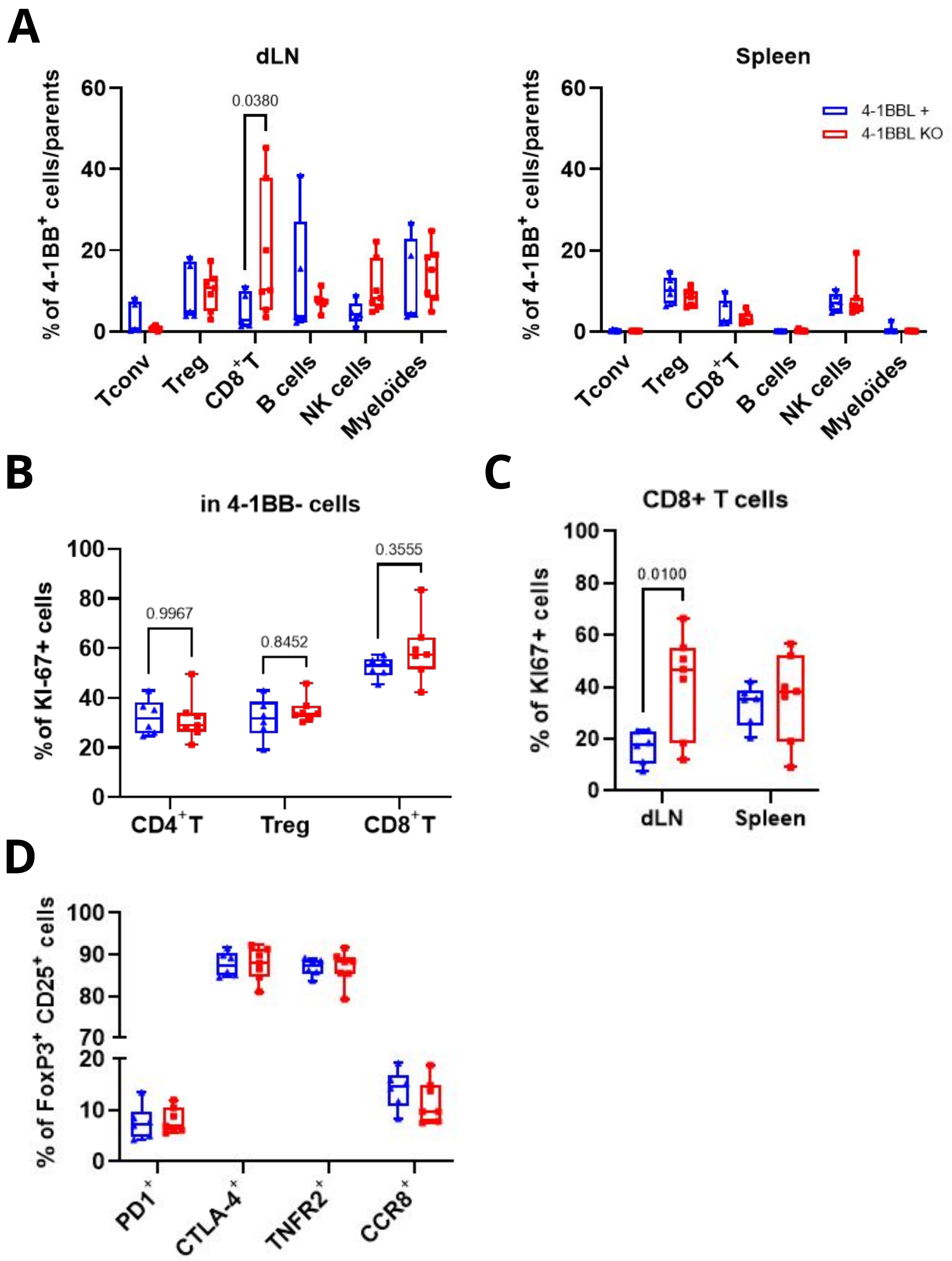

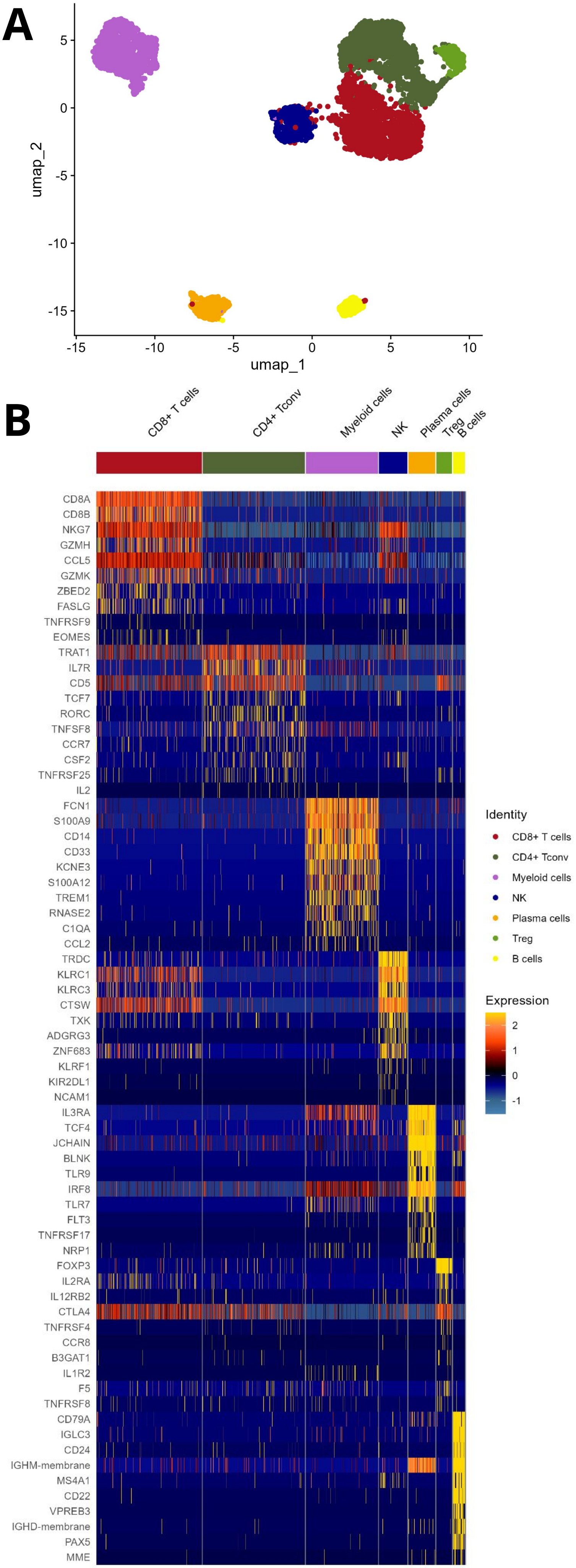

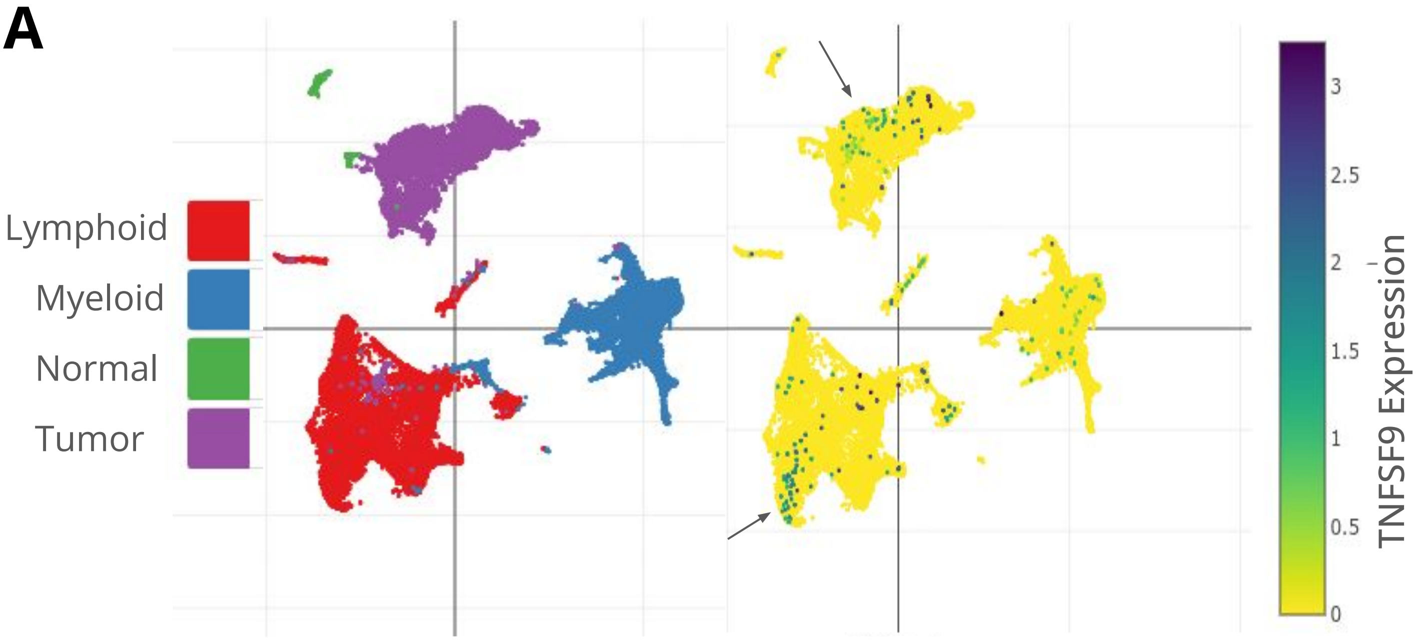

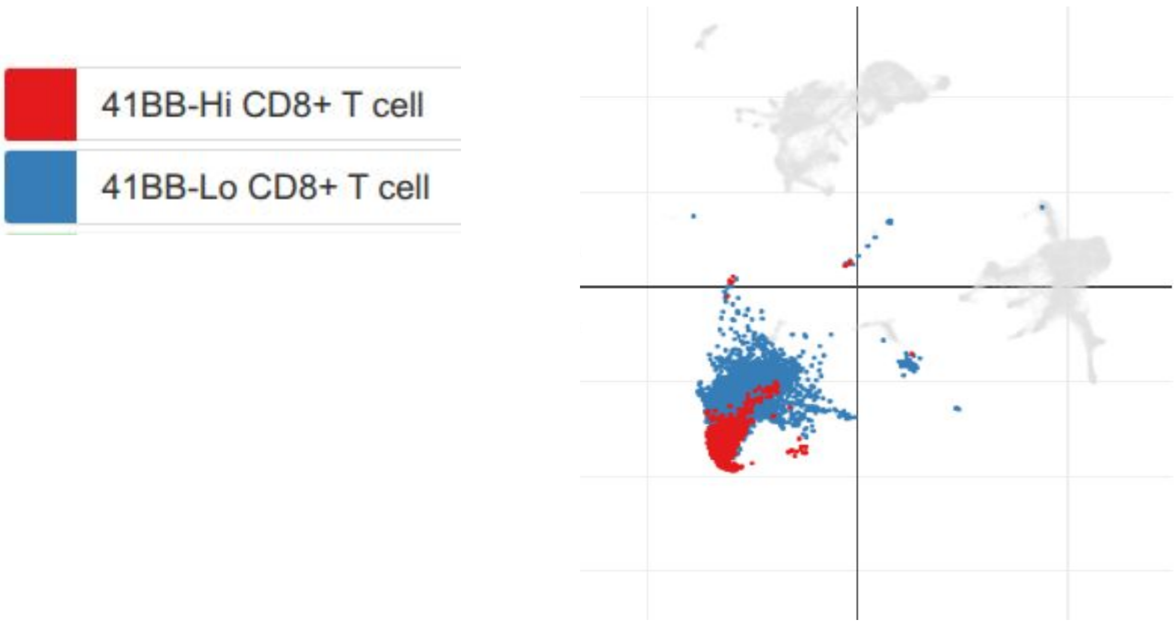
